# Closed-loop steering locates an evolutionary barrier to multidrug resistance in *Escherichia coli*

**DOI:** 10.64898/2026.09.17.752270

**Authors:** Atsushi Shibai, Hazuki Kotani, Chikara Furusawa

**Author notes:** **Corresponding author:** (AS), (CF).

## Abstract

Resistance to one drug often raises susceptibility to another; such collateral sensitivity is expected to constrain multidrug resistance. This is hard to test: failure to reach a state is indistinguishable from never having been driven toward it. We therefore recorded the change we imposed: a closed-loop system updated each lineage’s drug environment daily based on its measured distance to a target. Steered toward triple chloramphenicol–norfloxacin–kanamycin resistance for 70 days, 52 *Escherichia coli* lineages gained substantial resistance without converging on the target. The mismatch between imposed and observed movement mapped accessibility, locating a region immediately before the target where progress was reproducibly slowed. Crossing it was associated not with broad resistance mutations shared by most lineages, but with recurrent mutations upstream of the efflux pump gene *cmr*, which raised chloramphenicol resistance threefold without detectable cost. Progression toward multidrug resistance was therefore constrained without being prevented, at a location consistent with a known physiological trade-off; recording the imposed change makes accessibility measurable during evolution.

## Introduction

Resistance to different drugs cannot always be increased independently. This constraint is one rationale for combination therapy and drug cycling. The non-independence has a physiological basis: resistance to one drug is often accompanied by increased susceptibility to another, a relationship termed collateral sensitivity (Pál et al. 2015; Roemhild & Andersson 2021). For example, resistance to aminoglycosides can increase susceptibility to non-aminoglycoside drugs by reducing the proton motive force, which drives aminoglycoside uptake but also powers efflux pumps that expel other classes of antibiotics (Taber et al. 1987; Lázár et al. 2013). Such trade-offs have been systematically mapped (Imamovic & Sommer 2013; Lázár et al. 2014; Suzuki et al. 2014; Podnecky et al. 2018; Maeda et al. 2020) and used to design cycling schedules that can constrain the evolutionary paths toward multidrug resistance (Kim et al. 2014).

Whether such trade-offs actually impede progression toward multidrug resistance is a separate question. Collateral sensitivity is usually measured between endpoints, by assaying a population evolved under one drug against another. What matters instead is how a population moves during evolution when it is being pushed to raise several resistances at the same time, not which phenotypes it finally reaches. Populations may also circumvent a physiological conflict, and the outcome depends on history and genetic background: prior adaptation alters subsequent adaptation (Yen & Papin 2017), and the collateral response to a drug pair can be either sensitivity or cross-resistance (Nichol et al. 2019; Barbosa et al. 2019; Maltas & Wood 2019). More generally, constraints channel resistance evolution along a few directions (Maeda et al. 2020; Pinheiro et al. 2021; Iwasawa et al. 2022). What is not known is where, during progression toward a defined multidrug-resistant state, such constraints act, and what allows a population to overcome them.

Conventional experimental evolution is not well suited to answering this question. A drug condition is imposed and the resulting evolution observed (Baym et al. 2016), but the condition is not expressed as a direction and magnitude of change in resistance, and is not updated as resistance rises. In this situation, the failure of a population to reach a given state is ambiguous: the state may be inaccessible, or the imposed condition may simply not have pushed the population toward it. Feedback control solves part of this problem. The morbidostat raises the drug concentration as resistance evolves, so that selection does not relax (Toprak et al. 2012, 2013), and schedules designed in advance can manipulate and reverse resistance evolution (Nichol et al. 2015; Yoshida et al. 2017). However, the set point of a morbidostat is a level of growth inhibition rather than a state to be reached. It therefore fixes how strongly a population is challenged, but not the direction in which its environment is moved. A schedule that is fixed in advance also cannot respond to the current state of the population.

We therefore treated the imposed environmental change as an explicit, recorded input. We represented each population’s resistance as a vector in a three-dimensional space defined by resistance to three drugs, specified the target as a point in this space, and computed each passage’s drug environment from the difference between the measured resistance and the target. The environment was therefore updated daily and always directed at the target, and the magnitude of the update did not decrease as a lineage slowed, so a population that stopped advancing was still being pushed toward it.

Because both the imposed change and the resulting change in resistance are known at each step, the part of the observed movement that the imposed change does not account for can be extracted. We call this remainder the residual: it is small where a population followed the imposed change, and large where the movement was reduced or deflected. Residuals collected from many populations therefore describe how accessible each direction of movement was at each position in resistance space, and we summarised them as a scalar landscape whose downhill direction follows the residuals. Unlike a fitness landscape (Weinreich et al. 2006; de Visser & Krug 2014), its height measures not the resistance attained but the extent to which movement in the imposed direction was reduced or deflected. We call it an empirical accessibility landscape and use it to locate regions of reduced progression toward the target, which we term evolutionary barriers.

Feedback control has been proposed as a way of steering biological systems and managing pathogens (Lässig et al. 2023). Here it serves a further purpose: a population’s response to a change in its environment cannot be separated from its starting point and the path it happened to take unless that change is imposed and recorded, so control is also a way of measuring accessibility.

We applied this approach to chloramphenicol (CP), norfloxacin (NFLX) and kanamycin (KM), whose resistance mechanisms are expected to impose conflicting demands on membrane energetics. We evolved 52 independent lineages of a hypermutator *E. coli* strain (MDS42 Δ*mutS*) for 70 days under seven target schedules that directed lineages toward a common triple-resistance target from different directions during the final 30 days.

The reconstructed landscape located a reduced-accessibility region immediately before the target. Crossing was associated with recurrent mutations upstream of the efflux pump gene *cmr* (*mdfA*), rather than with broad resistance-associated mutations common to crossing and non-crossing lineages. These results locate where progression toward multidrug resistance is impeded, at a position consistent with a known physiological trade-off, and show that crossing is associated with an apparently regulatory change that adds resistance along a single constrained axis at no detectable cost.

## Results

### Closed-loop steering generates distinct approaches to triple resistance

We used target-directed closed-loop evolution to generate controlled approaches to a common multidrug-resistant state from different evolutionary histories. We defined a three-dimensional resistance space whose axes correspond to CP, NFLX and KM resistance (Fig. 1a). At each daily passage, the state of each evolving lineage was represented as a resistance vector, **R** = (*R*_CP_, *R*_NFLX_, *R*_KM_), where each component was the log_2_-transformed sub-MIC: the highest concentration that still supported a 100-fold increase in cell density over 24 h, read from a twofold dilution series (Methods).

**Figure 1.**
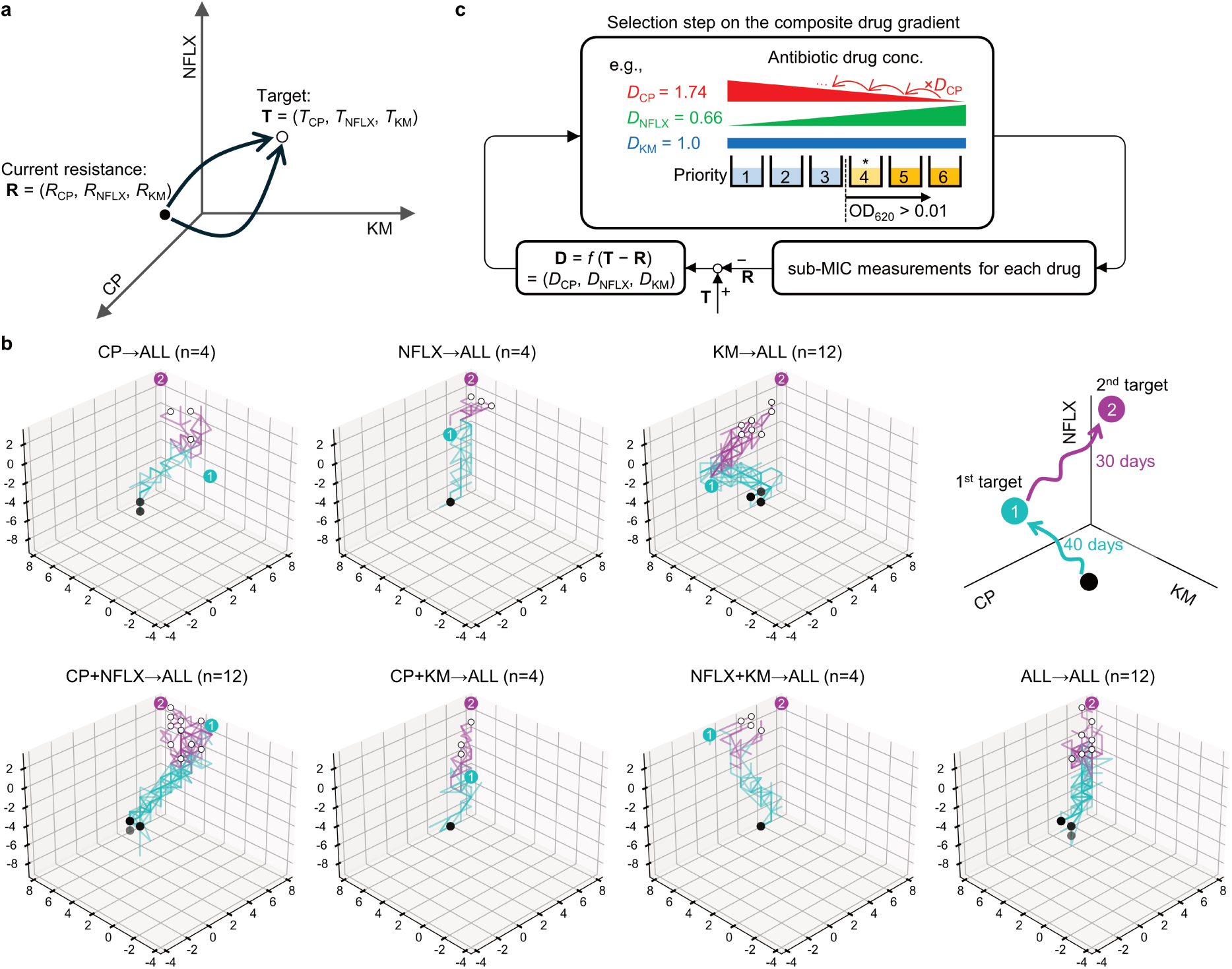
Closed-loop platform for steering bacterial evolution in three-drug resistance space. **a**, Schematic of the three-dimensional resistance space defined by chloramphenicol (CP), norfloxacin (NFLX) and kanamycin (KM). Each lineage has a current resistance state **R** = (*R*_CP_, *R*_NFLX_, *R*_KM_) and is driven toward a specified target **T** = (*T*_CP_, *T*_NFLX_, *T*_KM_). **b**, Seven target schedules and representative trajectories in log_2_ sub-MIC space. Cyan, the first 40 days toward assigned first-phase targets (label ①); magenta, the subsequent 30 days toward the common triple-resistance target ALL (label ②). In ALL→ALL, the two targets coincide; thus, only the second is shown. Black spheres indicate the ancestral starting point (day 1), and white spheres indicate the day-70 endpoints. The schematic at the top right shows the two-phase design and the orientation of the CP, NFLX and KM axes, which is the same in every subpanel and as in a; ticks show sub-MIC (log_2_ µg mL^−1^). **c**, At each 24 h cycle, cells were inoculated into a composite multidrug gradient. The panel shows the six dynamically updated gradient wells; the two lower-concentration rescue wells and the drug-free backup well (Methods) are omitted for clarity. Example dilution factors are shown (*D*_CP_ = 1.74, *D*_NFLX_ = 0.66, *D*_KM_ = 1.0): concentrations change by a factor *D*_j_ per step, so that CP rises and NFLX falls toward well 1 while KM is unchanged. The dilution factor **D** = (*D*_CP_, *D*_NFLX_, *D*_KM_) is updated according to the difference between the target and the current resistance, *D*_j_ = 2*^η^*^(*T*j−*R*j)^, for each drug *j*, where *η* = 1/‖**T** − **R**‖ when ‖**T** − **R**‖ > 1 and *η* = 1 otherwise, constraining *D*_j_ ∈ [2^−1^, 2^+1^]. Here *R*_j_ is obtained as the log_2_ sub-MIC from parallel single-drug twofold dilution assays, which isolate per-drug resistance from combination effects. Wells in the composite gradient are ranked by proximity to the target; among wells exceeding a growth threshold (OD_620_ > 0.01), the highest-priority well (asterisk) is propagated to seed the next cycle. Source data are provided with this paper.

We then defined target states as coordinate vectors in this resistance space. For each drug *j* ∈ {CP, NFLX, KM}, *R*^High^_*j*_ denotes the high-resistance target level, whereas *R*^Anc^_*j*_ denotes the resistance level measured for the ancestral strain. Each assigned target vector, **T** = (*T*_CP_, *T*_NFLX_, *T*_KM_), was specified by setting each component *T_j_* to either *R*^High^_*j*_ or *R*^Anc^_*j*_. Depending on whether one, two or three components are set to *R*^High^_*j*_, a target vector specifies single-drug, two-drug or triple resistance, giving seven distinct targets in total, excluding the case in which all three components are set to the ancestral level. For readability, we name each target by the drugs set to the high-resistance level: CP, NFLX, KM, CP+NFLX, CP+KM, NFLX+KM and ALL. Thus CP denotes (*R*^High^_CP_, *R*^Anc^_NFLX_, *R*^Anc^_KM_), CP+NFLX denotes (*R*^High^_CP_, *R*^High^_NFLX_, *R*^Anc^_KM_), and ALL denotes the triple-resistance target **T**_tri_ = (*R*^High^_CP_, *R*^High^_NFLX_, *R*^High^_KM_). This formulation allowed us to treat evolution as movement through this three-dimensional resistance space toward defined target coordinates.

To generate different approach histories, we implemented seven target schedules, each defined by a target vector **T**(*t*) that took one value during a first phase and another during a second (Fig. 1b). In the first phase, each lineage was steered for 40 days toward its assigned first-phase target; in the second phase, every lineage was redirected for a further 30 days toward the triple-resistance target ALL. We denote these schedules X→ALL, where X is the first-phase target. In ALL→ALL the two phases shared the same target, so those lineages were steered toward ALL throughout the 70-day experiment. The seven schedules were therefore designed to produce seven distinct approach histories to the same target coordinate.

We implemented these target schedules using a closed-loop culture system in which each lineage’s measured resistance vector was used to update the multidrug environment for the next passage (Fig. 1c). At each 24 h passage, the resistance vector **R** = (*R*_Cℓ_, *R*_NFLX_, *R*_KM_) was measured by parallel single-drug twofold dilution assays, which isolated per-drug resistance from combination effects. The difference between the assigned target **T** and the current state **R** was converted into drug-specific dilution factors, **D** = (*D*_Cℓ_, *D*_NFLX_, *D*_KM_), each a scaled exponential function of the corresponding component of **T** − **R** (Methods). The three drug-specific dilution series were then overlaid well by well, producing a single composite gradient of CP, NFLX and KM that stepped away from the lineage’s current operating point in the direction specified by the vector of log_2_ dilution factors. Among the wells in which the lineage grew, cells were propagated from the one closest to the assigned target. The controller thus set the direction of the concentration update, whereas survival determined how far along that direction each lineage was carried. Resistance measurement, target-directed environmental updating and survival-based selection were therefore coupled in a daily feedback loop.

Because the vector of log_2_ dilution factors was scaled to unit length along the direction of **T** − **R** (Methods), the controller applied a step of the same size toward the target at every passage, however far the target remained. A lineage that stopped advancing therefore continued to be pushed toward the target just as strongly. What was held constant was the magnitude of the nominal log_2_ dilution-factor vector, rather than the realised change in the concentrations selected for propagation.

We first confirmed that this control scheme could track resistance increases in single-drug evolution experiments, where the dilution range shifted upward over time (Supplementary Fig. 1). We next asked whether updating the direction of the environment each day, rather than fixing the direction of the update at the outset, was necessary. Using the three single-axis targets, we compared three schemes over 40 days (n = 12 lineages per scheme, pooled across the three targets). Under dynamic three-drug control, the direction of the concentration update was recomputed at every passage from the measured resistance, as described above. Under fixed three-drug control, this direction was set once at the start of the experiment and then held, while the concentrations themselves still followed growth; the two three-drug schemes therefore differed only in whether the direction of the concentration update was re-aimed as the population moved. Single-drug evolution under the corresponding drug alone was included as a reference, because evolution under the single corresponding drug is expected to be the easiest condition under which to approach a single-axis target.

The control scheme significantly affected the day-40 distance to the assigned target (two-way analysis of variance blocking on target, *P* = 0.0078). Fixed-direction control reached states farther from the target than either single-drug evolution (+1.07 log_2_ units, q = 0.017) or dynamic control (+0.92, q = 0.017), whereas dynamic control and single-drug evolution did not differ detectably (+0.15, q = 0.61) (Supplementary Fig. 2). Dynamic updating therefore approached the single-axis targets as closely as single-drug evolution did, whereas fixing the update direction degraded target approach, indicating that re-aiming the environment as resistance changes is necessary for the imposed gradient to remain effective.

We then applied this scheme to 52 independent lineages for 70 days under the seven target schedules. By day 40, the first phase of steering generated distinct approach histories in resistance space. Across 28 first-phase lineages, four from each schedule so that the seven schedules were equally represented, the assigned target was the nearest of the seven candidate targets far more often than expected by chance (mean rank of the assigned target 1.88 versus 4.0 expected; assigned target nearest for 18 of 28 lineages; exact schedule-to-target permutation test, *P* = 3.97 × 10⁻⁴), and in five of the seven schedules the day-40 states were significantly closer to their assigned target than to the other targets (per-schedule exact test, false discovery rate (FDR) < 0.05; Supplementary Fig. 3). The two exceptions were CP+KM and ALL, the only two targets requiring simultaneous increases along both the CP and KM axes. This weaker separation is consistent with previously reported collateral sensitivity between CP and KM (Lázár et al. 2013), and with conflicting selective demands along these two axes. Thus, the closed-loop system generated controlled trajectories in resistance space, and at the same time indicated that some target directions were harder to follow than others.

### Many trajectories approach but remain short of the triple-resistance target

Despite continuous target-directed updating of the environment, evolutionary trajectories did not converge uniformly on the ALL target. By day 70, many lineages had acquired substantial resistance increases, showing that the populations continued to adapt. However, their final resistance vectors were broadly distributed and often fell short of the target. In log_2_ sub-MIC coordinates, the across-lineage mean day-70 state, **R̅**_70_ = (6.0 ± 1.2, 0.9 ± 1.5, 6.3 ± 1.5) (± s.d.), lay below **T**_tri_ = (8.0, 3.0, 8.0) on all three axes. Euclidean distances in this space from the day-70 states to **T**_tri_ varied widely across lineages (median 4.1 log_2_ units; range 1.0–5.7).

Control operated on log_2_ sub-MIC because it was available within the daily cycle and identified the well from which to propagate, but because it is a threshold on a twofold series it changes only in whole steps and is a coarse descriptor of day-to-day movement. For the analyses below we instead fitted the full dose–response curves from the same assays to obtain log_2_ IC_50_ values, which are continuous. The same trajectories plotted in three-dimensional log_2_ IC_50_ space showed the same incomplete convergence toward the ALL target (Supplementary Fig. 4). We then projected the normalised log_2_ IC_50_ trajectories into a two-dimensional principal component analysis (PCA) space fitted across all lineages and time points. PC1 primarily captured CP and NFLX resistance and PC2 primarily captured KM resistance, explaining 58% and 32% of the variance, respectively.

We next asked whether the day-70 endpoints retained any imprint of the first-phase schedule. Endpoint position depended significantly on that schedule, both in the three-dimensional log_2_ IC_50_ space and in the PCA plane (permutational multivariate analysis of variance on Euclidean distances, PERMANOVA; R² = 0.68, *P* = 10^−4^ in both spaces; Anderson 2001). The spread of endpoints around each schedule mean did not differ among schedules (PERMDISP, the companion permutational test of homogeneity of multivariate dispersion; *P* = 0.42 and 0.12 in the three-dimensional space and the PCA plane, respectively; Anderson 2006), so the schedules differed in where their endpoints lay rather than in how widely these were scattered. The day-70 distance to the ALL target, however, did not differ significantly among schedules (Kruskal–Wallis, *P* = 0.17 in the three-dimensional space and *P* = 0.06 in the PCA plane, although the latter is close to the conventional threshold). Endpoint position therefore retained an imprint of the approach history, a form of historical contingency (Blount et al. 2018; Card et al. 2019), without any one schedule coming closer to the target than the others.

Applying the same drug-wise scaling and PCA loading matrix to **T**_tri_ gave its image in this plane, which we denote **x**_tri_ (Methods). The ancestral strain occupied a low-resistance region of the plane, whereas **x**_tri_ lay toward the high-resistance region (Fig. 2a). Lineages approached it to varying extents without reaching it. Steering nonetheless moved nearly all lineages toward **x**_tri_: the day-70 state lay closer to it than the day-40 state in 51 of 52 lineages, with a median reduction in distance of 34%. The single exception, an ALL→ALL lineage, is considered below. This motivated a quantitative analysis of where trajectory movement was reduced or deflected.

**Figure 2.**
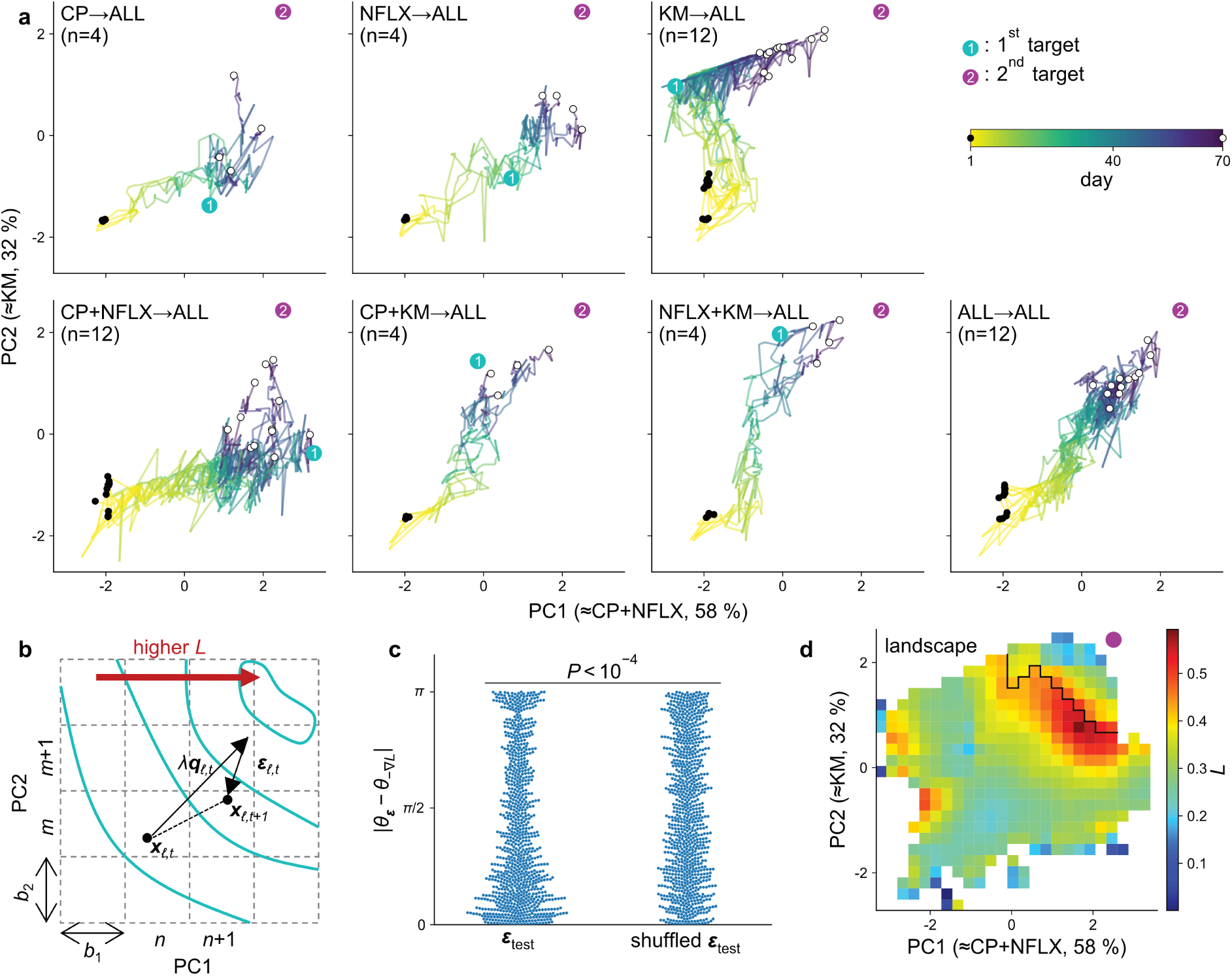
Reconstructed empirical accessibility landscape in PCA-transformed log_2_ IC_50_ space. **a**, Evolutionary trajectories under the seven target schedules (X→ALL); colours indicate time (day 1 to 70; colour bar). n is the number of independent lineages per schedule. Cyan ① and magenta ② mark the assigned first-phase target and the common triple-resistance target (ALL), projected into this plane. In ALL→ALL, the two targets coincide; thus, only the second is shown. Black dots mark the ancestral starting point (day 1), and white dots mark the day-70 final positions. Axes are principal components (PC1 ≃ CP+NFLX; PC2 ≃ KM) of normalised log_2_ IC_50_. **b**, Schematic of variables used for landscape reconstruction. For each lineage *ℓ* and day *t*, a lineage moved from **x***_ℓ_*_,*t*_ to **x***_ℓ_*_,*t*+1_ in the PCA-transformed log_2_ IC_50_ space, giving the realised displacement Δ**x***_ℓ_*_,*t*_ = **x***_ℓ_*_,*t*+1_ − **x***_ℓ_*_,*t*_. The target-directed dilution update generated a nominal control input from the log_2_ dilution factors, which was projected onto the PCA plane and denoted **q***_ℓ_*_,*t*_. This vector was scaled by a global factor *λ*, and the residual vector was defined as ***ε****_ℓ_*_,*t*_ = Δ**x***_ℓ_*_,*t*_ − *λ***q***_ℓ_*_,*t*_. Dashed lines show the grid, with cell indices (n, m) and widths b₁ and b₂, on which the landscape was represented and its gradient evaluated by forward finite differences; cyan curves are contours of the fitted landscape, which increases in the direction of the red arrow. The scalar landscape *L*(**x**) was fitted so that the landscape-derived direction −∇*L*(**x***_ℓ_*_,*t*_) aligned with ***ε****_ℓ_*_,*t*_. **c**, Angle concordance between observed and predicted residuals. Swarm plots show the absolute angular difference |*θ****_ε_*** − *θ*_−∇*L*_|, where *θ***_v_** denotes the direction of a vector **v** in the PCA plane and the difference is wrapped to [0, π], between the held-out residual vector ***ε****_ℓ_*_,*t*_ and the landscape-derived direction −∇*L*(**x***_ℓ_*_,*t*_). Left, held-out test data from 13 lineages, corresponding to 897 lineage-day intervals; right, the permutation whose test statistic was the median of 9,999 permutations, shown for reference. The observed angular differences were smaller than expected under permutation (one-sided permutation test, B = 9,999, *P* < 10⁻⁴; Methods). **d**, Reconstructed landscape on the PCA plane after Gaussian smoothing (h = 0.32; Methods); warmer colours denote higher *L*, and grid cells not visited by the evolving lineages are masked and shown in white (Methods). The black line marks a ridge in *L*, the operationally defined reduced-accessibility region (evolutionary barrier), which lies immediately before the projected triple-resistance target (magenta circle; see Methods). The landscape is shown as a descriptive summary; its detailed shape depends on the global gain. The residual field from which it is reconstructed was insensitive to that gain over a fourfold range around the value used (Supplementary Fig. 5e), and the ridge shown here was returned unchanged at twice that value, but no closed boundary was obtained at half of it (Methods). Source data are provided with this paper.

### Residual trajectory analysis reveals a reduced-accessibility region near the triple-resistance target

To determine whether incomplete progression toward **x**_tri_ reflected a structured constraint rather than random lineage-to-lineage variation, we compared the controller-defined update with the phenotypic movement that actually occurred. For each lineage *_ℓ_* and day *t*, we denoted the position of the lineage before and after one 24 h passage in the PCA-transformed log_2_ IC_50_ space as **x***_ℓ_*_,*t*_ and **x***_ℓ_*_,*t*+1_, respectively, and defined the observed displacement as Δ**x***_ℓ_*_,*t*_ = **x***_ℓ_*_,*t*+1_ − **x***_ℓ_*_,*t*_. For the same transition, the concentration update specified by the controller, expressed as the vector of log_2_ dilution factors applied to the three drugs, was projected into the same plane using the same drug-wise scaling and PCA loading matrix, giving a two-dimensional control vector **q***_ℓ_*_,*t*_ (Methods). The controller specifies the direction and the relative magnitude of each concentration update, but not the phenotypic displacement that this update produces: the conversion from a log_2_-fold change in imposed drug concentration to movement in the PCA plane is not known a priori. We therefore introduced a single global gain, *λ*, defined as the ratio of the summed observed displacement lengths to the summed control-vector lengths, *λ* = ∑*_ℓ_*_,*t*_ǁΔ**x***_ℓ_*_,*t*_ǁ/∑*_ℓ_*_,*t*_ǁ**q***_ℓ_*_,*t*_ǁ, where both sums run over every lineage-day interval in the dataset (52 lineages × 69 daily transitions, N = 3,588), so that controller-defined inputs and observed displacements carry the same total magnitude overall. *λ***q***_ℓ_*_,*t*_ therefore serves as a reference for the displacement expected from the controller-defined update alone, and we defined the residual movement as ***ε****_ℓ_*_,*t*_ = Δ**x***_ℓ_*_,*t*_ − *λ***q***_ℓ_*_,*t*_ (Fig. 2b). Because *λ* is a single scalar estimated once across all 52 lineages, it sets only the overall scale of the reference displacement and cannot by itself create position-dependent structure. The sensitivity of the reconstruction to its value is examined below (Supplementary Fig. 5a–f).

These residuals provide an operational measure of how trajectories deviated from the movement expected from the controller-defined update. Spatially structured residuals would indicate regions where similar controller-defined inputs tend to produce reduced or deflected movement, whereas residuals arising from lineage-to-lineage variation alone would not align reproducibly with position in the PCA plane. We therefore fitted an empirical accessibility landscape *L*(**x**) in the PCA plane such that its local downhill direction, −∇*L*(**x**), aligned with the residual vectors. High regions of *L*(**x**) thus mark positions from which trajectories tended to be deflected or to fail to progress, whereas low regions correspond to more accessible directions of movement. *L*(**x**) was estimated on a regular grid in the PCA plane, with cells not visited by the evolving lineages left undefined. Details of the scaling, grid construction, masking criterion and numerical optimisation are provided in Methods. Its height therefore measures not the resistance a lineage has attained, but the extent to which the imposed, target-directed perturbation failed to produce the movement it specified.

We tested whether this inferred landscape captured reproducible structure rather than overfitting the observed trajectories. We split the 52 lineages into training and held-out test sets, reconstructed the landscape from the training lineages, and asked whether the inferred local field aligned with residual movements in the held-out lineages. The angular difference between observed residual vectors and predicted landscape-derived directions was significantly smaller than that obtained after randomly reassigning residual vectors among test positions (one-sided permutation test, 9,999 permutations, *P* < 10⁻⁴; Fig. 2c). Thus, the residual movements contained reproducible spatial structure that was not explained by the nominal control input alone.

The landscape reconstructed from all lineages revealed a ridge-like region immediately before **x**tri (Fig. 2d). Many trajectories approached this region during the 70-day experiment, but progression toward **x**_tri_ was reduced there, and only a subset of lineages crossed into the sparsely sampled target-side region beyond the ridge. The one lineage that did not end closer to **x**_tri_ than it had been at day 40 illustrates this: this ALL→ALL lineage had already reached the ridge by day 40 and subsequently fluctuated along it without crossing. Because that target-side region was reached by only a few lineages, we do not interpret the apparent decrease of *L* beyond the ridge; the ridge itself, however, was traversed by trajectories from multiple independent lineages and is an internal feature of the sampled landscape rather than an artefact of its masked boundary (Supplementary Fig. 5a, b). We use the term evolutionary barrier operationally to denote this reduced-accessibility region: it is not an absolute boundary that prevents triple resistance, but a region in which controlled trajectories were reproducibly slowed or deflected, or failed to progress toward **x**_tri_.

The reconstruction is not sensitive to the precise value of *λ*. Over a fourfold range around the moment-matched value (0.549), from *λ*/2 to 2*λ*, the per-cell residual directions agreed closely (cell-weighted mean cosine ≥ 0.60 for every pair; part of this agreement is expected, because the residual is increasingly dominated by the shared −*λ***q** term), whereas discarding the control input (*λ* = 0) gave an unrelated field (cosine 0.06 against *λ* and −0.16 against 2*λ*; Supplementary Fig. 5e). The ridge is not directly a property of that field, because the landscape was refitted at each gain, and it was robust over a narrower range: the detected boundary was identical cell for cell at *λ* and 2*λ*, whereas at *λ*/2 the detected cells left gaps too wide to bridge, so no target-side region is defined at that gain (Methods). Because the projected control vector points predominantly along the target direction, the scaled reference displacement *λ***q***_ℓ_*_,*t*_ and the fitted gradient ∇*L*(**x**) are close to parallel over most of the sampled plane, so the global gain and the landscape are only weakly separable: an increase in *λ* can be partly absorbed by an increase in the fitted height of *L*(**x**). We therefore treat the landscape as a descriptive summary, and define barrier crossing below by whether a lineage entered the target-side region, independently of the fitted height of *L*.

### Barrier crossing is associated with upstream *cmr* mutations

We therefore asked whether crossing the inferred barrier was associated with specific genetic changes, and in particular with a change that raises resistance along one axis without incurring the collateral cost that produces the trade-off. We performed whole-genome sequencing of evolved populations from the three schedules with the largest number of replicate lineages: KM→ALL, CP+NFLX→ALL and ALL→ALL. All 12 lineages from each schedule were sequenced at day 70, and the corresponding day-40 samples were also analysed to distinguish mutations that arose during the first-phase approach from those that emerged or were enriched during the subsequent 30 days of steering toward ALL. Using the barrier definition described in Methods (Supplementary Fig. 6), we classified lineages whose trajectories entered the target-side region at least once as Crossed, and those that remained on the pre-barrier side as Stalled. Among the 36 sequenced lineages, 14 were classified as Crossed and 22 as Stalled. Crossing was more frequent in KM→ALL (8/12) than in CP+NFLX→ALL (3/12) or ALL→ALL (3/12), though this difference was not significant (Freeman–Halton extension of Fisher’s exact test, *P* = 0.075).

Mutations accumulated at many loci across these lineages, as expected from the Δ*mutS* background. Recurrently mutated loci included genes associated with drug efflux, membrane permeability and antibiotic response, such as *acrAB*, which encodes components of a resistance-nodulation-division (RND) efflux system (Ma et al. 1995; Nikaido & Pagès 2012). These loci were mutated in both Crossed and Stalled lineages (Fig. 3). We therefore asked whether any locus was specifically enriched in Crossed lineages. Among loci mutated in at least five of the 36 sequenced lineages, we compared day-70 mutation frequencies between Crossed and Stalled groups using Fisher’s exact tests followed by false discovery rate correction. This analysis identified a single significantly enriched locus: the intergenic region between *ybjG* and *cmr*. Mutations in this region were present in 8 of 14 Crossed lineages but only 1 of 22 Stalled lineages (*P* = 7.2 × 10⁻⁴, FDR < 0.05; odds ratio 28, 95% confidence interval 3–131). By contrast, the broad resistance-associated loci that were mutated in most lineages did not distinguish the two groups: mutations in *acrB* (13 of 14 Crossed versus 21 of 22 Stalled), *acrR*, *marR*, *ompF* and *tolC* were common on both sides of the barrier and showed no significant enrichment (all *P* > 0.1).

**Figure 3.**
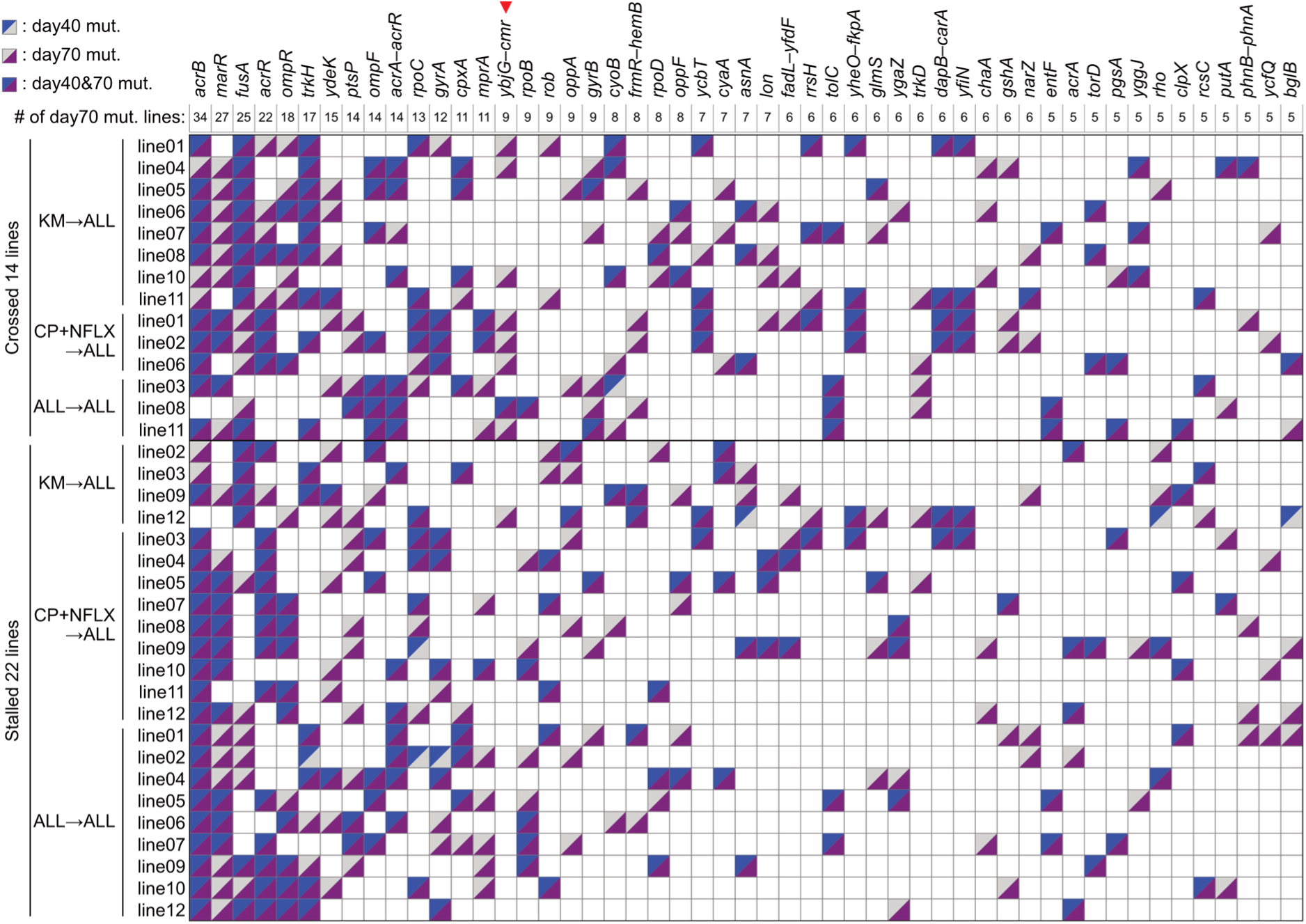
Distribution of mutations across evolved lineages and frequently mutated genes at days 40 and 70. Presence or absence of mutations is shown across the 36 sequenced evolved *E. coli* lineages (rows), comprising 12 lineages from each of the KM→ALL, CP+NFLX→ALL and ALL→ALL schedules, and genes or intergenic regions mutated in ≥5 lineages at day 70 (columns). Genes are ordered by the number of mutated lineages at day 70, as indicated below gene names. Lineages are grouped according to whether they crossed the evolutionary barrier (Crossed, n = 14) or stalled (Stalled, n = 22), as well as by target schedule (e.g., KM→ALL). Blue upper-left and purple lower-right triangles indicate mutations detected at day 40 and day 70, respectively; cells with both triangles indicate mutations detected at both time points. Notably, mutations in the *ybjG*–*cmr* intergenic region (red arrowhead above the column) were present in 8 of 14 Crossed but only 1 of 22 Stalled lineages (Fisher’s exact test, *P* = 7.2 × 10⁻⁴; FDR < 0.05) and were largely absent at day 40, suggesting a late-onset, barrier-associated mutational signature. Triangles indicate consensus genotype calls; read-level allele frequencies at day 40 for the *ybjG*–*cmr* positions are given in Supplementary Note 1. Broad resistance loci (e.g. *acrB*, mutated in 13 of 14 Crossed and 21 of 22 Stalled lineages) did not differ between groups (Fisher’s exact test, *P* > 0.1); only the *ybjG*–*cmr* intergenic region was significantly enriched (odds ratio 18.7). Background and experiment-wide variants were excluded before this analysis (Methods). Source data are provided with this paper.

We next examined when these mutations appeared. Because mutations arising in the first phase can reflect the first-phase target rather than the approach to ALL, we compared the *ybjG*–*cmr* region at day 40 and day 70. In the consensus genotypes these mutations were present in one of the 36 sequenced lineages at day 40 but in 9 by day 70 (Fig. 3); a read-level re-analysis of the day-40 alignments detected a variant in three of those nine lineages at day 40, at frequencies of 0.42 to 0.97 (Supplementary Note 1). They were therefore unlikely to be primary drivers of the initial target-specific resistance increases, and instead arose or increased in frequency during the later phase, when lineages had already approached the reduced-accessibility region near **x**_tri_.

The *ybjG*–*cmr* intergenic region lies between two divergently transcribed genes. *cmr*, also known as *mdfA*, encodes the major facilitator superfamily (MFS) multidrug efflux pump Cmr/MdfA, a proton antiporter that exports chloramphenicol and has an unusually broad substrate range (Edgar & Bibi 1997; Mine et al. 1998; Heng et al. 2015), whereas *ybjG* encodes a neighbouring predicted undecaprenyl-diphosphatase. The interval is upstream of both genes, but the mutations observed in evolved lineages were not distributed evenly across it. Instead, all fell within about 100 bp of the *cmr* start codon, near annotated promoter elements upstream of *cmr* (Fig. 4a). Hereafter we refer to these variants as upstream *cmr* variants.

**Figure 4.**
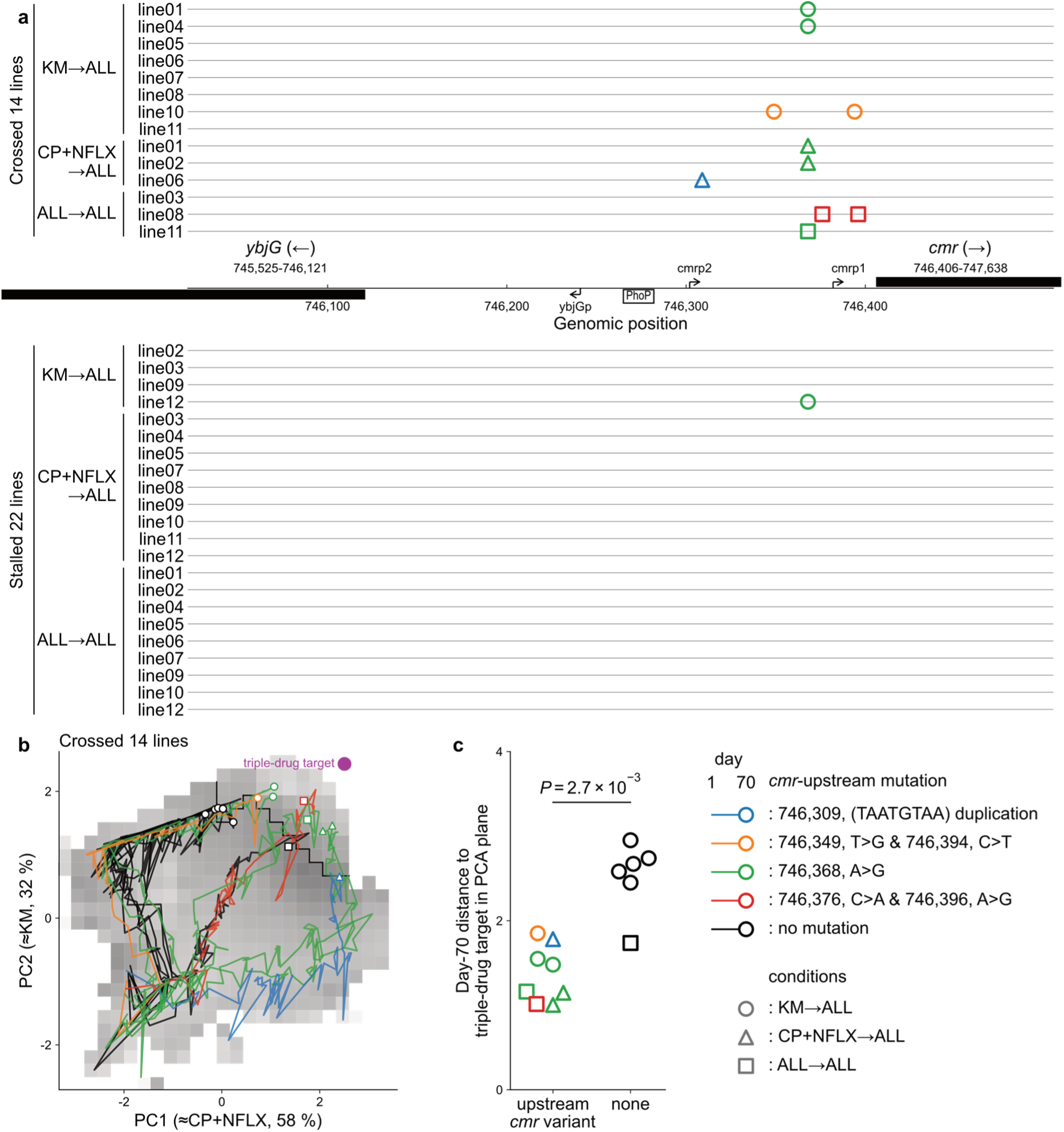
Mutations in the *ybjG–cmr* intergenic region are associated with barrier crossing and proximity to the triple-resistance target. a, Distribution of mutations identified within the *ybjG–cmr* intergenic region across the 36 sequenced evolved *E. coli* lineages. Rows represent individual lineages grouped by evolutionary outcome (Crossed or Stalled) and target schedule. Symbols denote the presence of specific mutations; symbol colour indicates the mutation type, with genomic coordinates and nucleotide substitutions given in the legend, and symbol shape indicates the target schedule. The highlighted mutations were all located upstream of *cmr*, near annotated regulatory elements, and were predominantly observed in Crossed lineages. Black bars show the coding regions of *ybjG* and *cmr*, with arrows indicating their transcriptional directions. Promoter annotations (*cmrp1*, *cmrp2* and *ybjGp*) and the PhoP binding site were curated from RegulonDB (Tierrafría et al. 2022). **b,** Evolutionary trajectories of the 14 Crossed lineages projected onto the accessibility landscape reconstructed from PCA-transformed log_2_ IC_50_ changes (see Fig. 2). The greyscale background indicates the inferred accessibility landscape *L*, with darker shading denoting higher *L*, and the black line indicates the estimated barrier. Each trajectory is coloured according to the mutation type shown in **a**; lineages without mutations in the *ybjG–cmr* intergenic region are shown in black. Symbols mark the day-70 endpoints of each trajectory. The magenta point indicates the projected triple-resistance target. **c,** Day-70 distances from the triple-resistance target in the PCA plane for Crossed lineages with or without upstream *cmr* mutations. Each point represents one lineage, with the outline colour corresponding to the mutation type shown in **a** and **b** and the marker shape indicating the target schedule. Crossed lineages carrying upstream *cmr* mutations reached positions closer to the triple-resistance target than those without such mutations. The *P* value was calculated using a two-sided Mann–Whitney U test. The legend at right applies to a–c. Source data are provided with this paper.

Across the 36 sequenced lineages, nine carried an upstream *cmr* variant, comprising six distinct mutations. These included single-nucleotide substitutions, a short repeat expansion and closely spaced substitutions. The most frequent was the 746,368 A>G substitution, which arose independently in six lineages; others included expansion of the TAATGTAA repeat at position 746,309 from one to two copies. None fell within the annotated regulatory elements themselves, the *ybjG* promoter and a PhoP binding site, although all lay close to the annotated *cmr* promoters. Both the positional bias and the recurrent, independent origin of these variants are consistent with positive selection acting on the regulatory region upstream of *cmr*. None was detected in four drug-free control lineages propagated in parallel for 70 days, suggesting that they arose under drug selection rather than as neutral variation in the mutator background.

Upstream *cmr* variants were also associated with where lineages ended on the reconstructed landscape. Among the 14 Crossed lineages, those carrying such a variant ended closer to **x**_tri_ at day 70 than those without (two-sided Mann–Whitney U test, *P* = 2.7 × 10⁻³; Fig. 4b,c). This association did not depend on the Crossed/Stalled classification or on the fitted height of *L*: across all 36 sequenced lineages, those carrying an upstream *cmr* variant ended closer to **x**_tri_ than those without (median distance 1.5 versus 2.5 units in the PCA plane; *P* = 1.3 × 10⁻³; Supplementary Fig. 7). This association makes no reference to the barrier, and is therefore independent of the fitted landscape and of the gain.

However, upstream *cmr* variants were neither necessary nor sufficient for barrier crossing. Six of the 14 Crossed lineages lacked detectable mutations in this intergenic region, and one lineage carrying the recurrent 746,368 A>G variant ended as far from **x**_tri_ as lineages without any such variant (Supplementary Fig. 7). This argues against a model in which an upstream *cmr* mutation acts as a causal switch. Instead, upstream *cmr* variants appear to be recurrent, background-dependent modifiers that are associated with further progress toward **x**_tri_ in some already adapted lineages.

### A recurrent upstream *cmr* mutation provides a CP-biased resistance gain

To test whether an upstream *cmr* mutation can directly alter resistance phenotypes, we introduced the most frequently observed variant, 746,368 A>G, into the parental strain and compared its resistance phenotypes with those of the parental strain and the Δ*mutS* control. The engineered strain showed an approximately threefold increase in CP resistance compared with the parental strain, whereas the Δ*mutS* control differed from the parental strain by less than one dilution step on every axis (Fig. 5a). Resistance to NFLX and KM was not detectably decreased under the same single-drug assay conditions, at a precision sufficient to have detected a decrease of less than 1.1-fold; both axes showed small increases instead (NFLX, +0.30; KM, +0.10 log_2_ units). The engineered strain also did not show a detectable reduction in growth rate in drug-free medium under the same culture conditions (Supplementary Fig. 8a). Thus, the upstream *cmr* mutation was sufficient, in the parental background, to provide a CP-biased resistance gain, although it did not reproduce the full multidrug-resistance phenotype of Crossed lineages.

**Figure 5.**
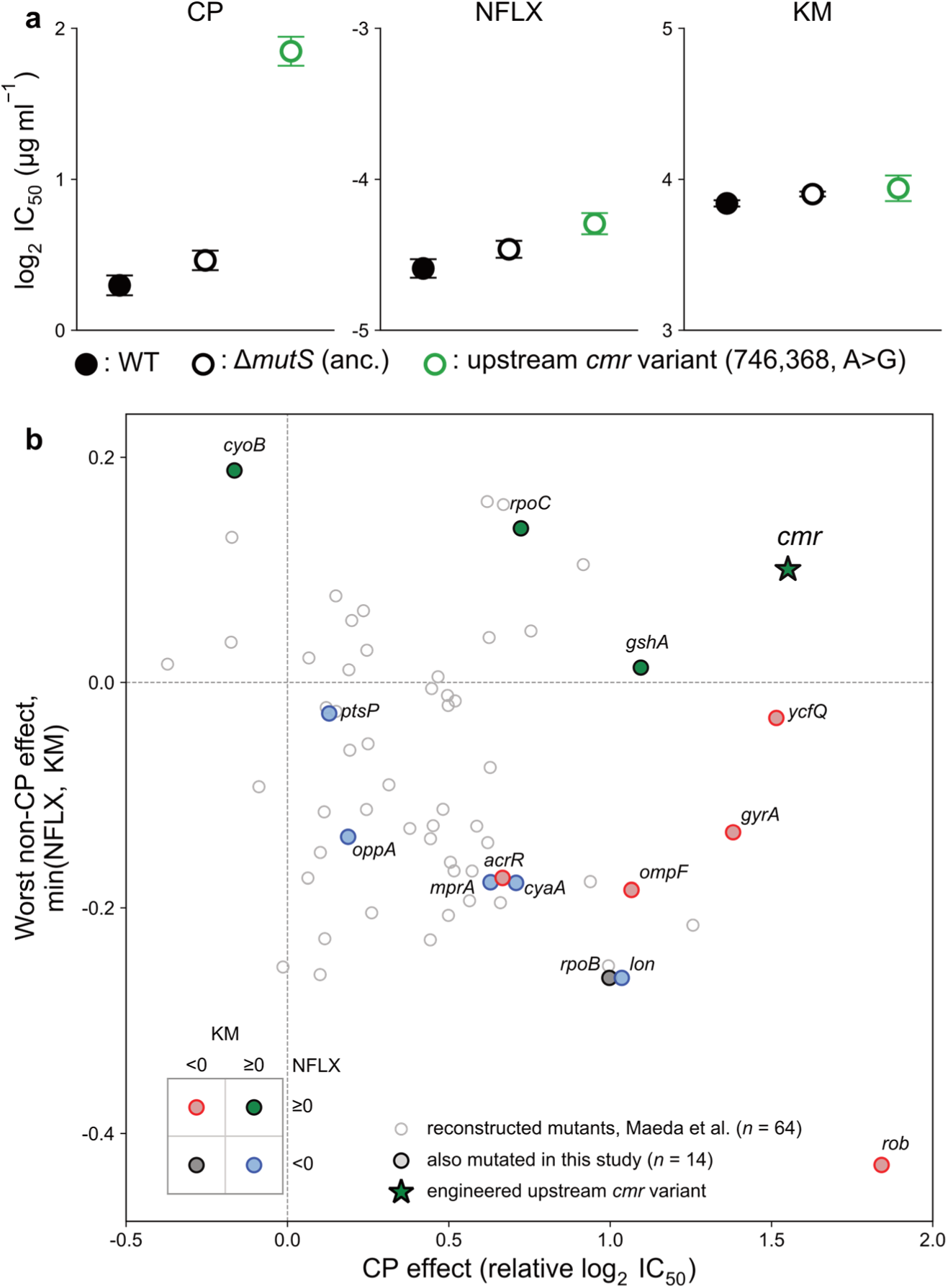
Upstream *cmr* mutation provides a CP-biased resistance gain without detectable cost along other drug axes. **a**, Resistance phenotypes of parental MDS42 (WT), MDS42 Δ*mutS* and the engineered MDS42 strain carrying the 746,368 A>G upstream *cmr* mutation. Introduction of the mutation increased resistance specifically to chloramphenicol (CP) by approximately threefold, with no detectable decrease along the norfloxacin (NFLX) or kanamycin (KM) axes, which instead showed small increases (+0.30 and +0.10 log_2_ units, respectively). Data are shown as mean ± s.d. (n = 8), with individual replicates indicated by dots. The assay was precise enough to exclude a meaningful cost along the other axes: with a replicate standard deviation of ≈ 0.06 log_2_ units on these axes (n = 8), a decrease of ≥ 0.1 log_2_ units, less than a 1.1-fold change, would have been detected on either the NFLX or KM axis at 80% statistical power (two-sided test, α = 0.05). The Δ*mutS* control differed from the parental strain by at most 0.17 log_2_ units on any axis, below one dilution step; although this difference was detectable at the precision of the assay (n = 8, *P* < 0.001), it is negligible relative to the several-log_2_-unit resistance changes acquired during evolution, which can therefore be attributed to acquired mutations rather than to the mutator background itself. **b**, Comparison of resistance changes caused by the engineered upstream *cmr* mutation and by reconstructed mutant strains from Maeda et al. (2020). The horizontal axis shows the CP effect and the vertical axis the worst non-CP effect, defined as the smaller of the NFLX and KM effects; all effects are expressed as log_2_ fold changes in IC_50_ relative to the corresponding parental strain, and dashed lines mark zero effect. The upstream *cmr* point (green star) was calculated from the resistance assay shown in a as a fold change relative to parental MDS42. Other points correspond to the 64 MAGE-reconstructed mutant strains from Maeda et al. (2020); the 14 in gene regions overlapping recurrently mutated loci in our evolved lineages are highlighted as larger symbols with gene labels. Symbol colour indicates the signs of the NFLX and KM effects, as shown in the key at lower left. The engineered upstream *cmr* mutation increased CP resistance without a detectable decrease in NFLX or KM resistance. Source data are provided with this paper.

We next asked whether this CP-biased effect was distinctive relative to the resistance effects of other reconstructed resistance-associated mutations. For comparison, we used the reconstructed mutant panel reported by Maeda et al. (2020), in which 64 representative mutations identified from high-throughput laboratory evolution were individually introduced into the parental strain used in that study by multiplex automated genome engineering (MAGE; Wang et al. 2009) and assayed for resistance to a panel of antibacterial compounds, including CP, NFLX and KM. We compared the CP, NFLX and KM resistance changes of all 64 reconstructed mutants with those of the engineered upstream *cmr* mutant, highlighting the 14 mutants in gene regions that overlapped with recurrently mutated loci identified in our evolved lineages (Fig. 3). Although upstream *cmr* mutations also arose in the Maeda et al. evolution experiments, they were not among these 64 reconstructed variants, so the comparison is with routes other than this one.

Many of the reconstructed mutants from Maeda et al. showed opposing effects across drug axes, consistent with trade-offs in this multidrug resistance space. By contrast, the engineered upstream *cmr* mutation combined a large CP gain with no detectable decrease in NFLX or KM resistance (Fig. 5b). Across the full panel, no reconstructed mutant matched this combination (CP gain ≥ 1.55 log_2_ units with no decrease along the NFLX or KM axes; 0 of 64; empirical *P* = 0.015). Its CP gain lay at the 98th percentile of the 64 mutants, and the single mutant with a larger CP gain incurred a decrease along another axis.

To ask whether these variants act through *cmr* expression, we re-analysed expression profiles from the Maeda et al. (2020) dataset. In each of the chloramphenicol and puromycin experiments, the single lineage carrying the 746,309 TAATGTAA repeat expansion, which was also recovered in our evolved lineages although it was not the variant we reconstructed, showed approximately eightfold higher *cmr* expression than the three lineages lacking it (Supplementary Fig. 8b). Because this variant differs from the one whose resistance effect we measured and arose in a different evolutionary context, this comparison is descriptive. It is nonetheless consistent with at least some upstream *cmr* variants increasing *cmr* expression by altering the regulatory region upstream of the gene.

The engineered 746,368 A>G variant raised CP resistance by about 1.55 log_2_ units, a modest gain relative to the several log_2_ units acquired along that axis over 70 days. Together with the trajectory-level results, the genetic, phenotypic and expression data are consistent with upstream *cmr* variants acting as secondary modifiers on already adapted genetic backgrounds, providing a locally favourable increment of CP resistance in lineages that have approached the constrained region.

## Discussion

We set out to distinguish a genuine constraint on the evolution of multidrug resistance from a mere failure to evolve toward that state. We therefore treated the imposed environmental change as an explicit, recorded quantity: each lineage’s resistance was measured daily and used to compute its next drug environment from the difference between the measured state and a defined target, so that the direction and magnitude of the controller-defined update were known at every step. Because the update was scaled to unit length, its magnitude did not diminish as a lineage slowed. What was recorded was this environmental input, not the realised strength of selection, which depends on genetic background and physiological state. Comparing this recorded update with the phenotypic movement that followed provided a local measure of how the resistance space responded to directed perturbation. Applied to 52 lineages steered toward simultaneous resistance to CP, NFLX and KM, the approach revealed a reduced-accessibility region immediately before the triple-resistance target, which only a subset of lineages crossed, and identified recurrent mutations upstream of *cmr* (*mdfA*) as the only locus significantly enriched among the lineages that crossed it.

The quantity on which this measurement rests is the residual: the part of the observed movement that the controller-defined update does not account for. Because the residual is defined relative to a recorded input, it carries information that trajectories alone do not. A landscape reconstructed from observed trajectories, with resistance itself as its altitude (Iwasawa et al. 2022), is validated by the observation that populations ascend it; within such a construction, a population that stops moving lies at a peak, and a barrier cannot be defined because the driving is not measured independently. The present reconstruction inverts that relationship: control is not designed from a landscape but used as the instrument by which a landscape is measured.

The residuals contained spatial structure that was reproducible in held-out lineages, and several alternative explanations for this structure can be excluded. First, the reduced-accessibility region does not coincide with cells in which the projected control input was systematically weaker or stronger (Supplementary Fig. 5c,d), so it is not an artefact of uneven driving. Second, the region lies in the densely sampled interior of the plane rather than at its masked boundary (Supplementary Fig. 5a,b), so it is not an edge effect of the mask. Third, the same structure is recovered from the day-40 to day-70 interval alone (Supplementary Fig. 5f), so it is not generated by the first-phase portion of the trajectories, during which lineages were still approaching different targets. The structure is also unlikely to reflect a uniform decline in responsiveness at higher resistance, of the kind expected from diminishing-returns epistasis (Chou et al. 2011; Kryazhimskiy et al. 2014), because such a decline would not produce a localised ridge. The fitted height and orientation of the landscape depend on the global gain, which is only weakly separable from the landscape itself; the residual field and the ridge-derived boundary were nonetheless stable over the range of gains examined (Supplementary Fig. 5e).

The location of the reduced-accessibility region is consistent with a physiological conflict that has already been documented, and our contribution is to locate this conflict rather than to discover it. Uptake of aminoglycosides such as KM is driven by the proton motive force (PMF) (Bryan & Kwan 1983; Taber et al. 1987; Allison et al. 2011), so KM resistance can be gained by lowering PMF, whereas efflux of CP and NFLX is mediated by proton antiporters that depend on the same force (Paulsen et al. 1996; Nikaido & Pagès 2012). The triple-resistance region can therefore be entered only by moving along both PCA axes at the same time, raising CP/NFLX and KM resistance together, which is precisely where competing demands on membrane energetics are expected to arise. Consistent with this, the two first-phase schedules whose day-40 states separated least from the other targets were the two requiring simultaneous increases along both the CP and KM axes (Supplementary Fig. 3). What the closed-loop measurement adds is where, during an active approach to triple resistance, this conflict impedes progress. We did not measure PMF, membrane potential, efflux activity or drug uptake, so this physiological interpretation remains a hypothesis.

If the region reflects such a conflict, the kind of mutation that can carry a lineage past it is restricted. A further general increase in resistance would draw on the same contested resource, the PMF. What is required instead is an increase along one axis without imposing a detectable collateral cost under the assay conditions. The sequencing results matched this expectation. Mutations broadly associated with resistance, including those affecting the RND-type AcrAB-TolC system, were common on both sides of the region and did not distinguish crossing from non-crossing lineages. What distinguished the two groups were recurrent mutations upstream of *cmr* (*mdfA*), which encodes an MFS multidrug efflux pump (Edgar & Bibi 1997; Mine et al. 1998). Because this pump is itself driven by the proton motive force, we cannot argue from its mechanism that raising *cmr* expression spares the contested resource. The evidence that this route is inexpensive is instead empirical. Introduced into the unevolved parental strain, the reconstructed 746,368 A>G variant raised CP resistance approximately threefold with no detectable decrease along the NFLX or KM axes. The variant also caused no detectable reduction in growth rate in drug-free medium. Within the resolution of these assays, the variant therefore adds CP resistance at little or no cost. Across the 64 reconstructed mutants of Maeda et al. (2020), none achieved a comparable CP gain while maintaining resistance along both of the other axes. In a re-analysis of the same dataset, a second upstream variant, the 746,309 repeat expansion that also arose in our evolved lineages, was associated with an approximately eightfold increase in *cmr* expression, which is consistent with a regulatory mechanism.

These variants nonetheless act as conditional modifiers rather than as switches. They were largely absent at day 40 and frequent at day 70, and were enriched among crossing lineages and associated with more target-proximal endpoints. This last association held across all sequenced lineages, independently of the barrier classification. Yet the variants were neither necessary nor sufficient: several crossing lineages lacked them, and one lineage carrying the recurrent variant ended as far from the target as lineages without it. Moreover, the threefold CP gain is modest relative to the resistance acquired over 70 days. Crossing is therefore unlikely to be governed by the waiting time for a single rare mutation, particularly in a hypermutator background in which mutational supply is not limiting. It depends instead on the genetic background on which such a variant acts: a small, apparently cost-free increase becomes useful only after a lineage has otherwise approached the constrained region. This suggests a more general possibility. In regions of phenotype space that are governed by physiological trade-offs, the change that allows progress may often be a narrow regulatory adjustment rather than one of the broad resistance mechanisms that are common elsewhere: an adjustment that adds capacity along one axis without imposing a detectable collateral cost under the assay conditions.

Several features of the design bound these conclusions. The Δ*mutS* background elevates the point-mutation rate by roughly two orders of magnitude, as measured for mismatch-repair-deficient *E. coli* (Lee et al. 2012), and was chosen so that the supply of resistance mutations would not itself limit the response to steering. Adaptation is therefore selection-limited rather than mutation-limited, which strengthens the inference that a region in which lineages stall despite abundant mutational supply reflects the structure of accessibility rather than the waiting time for beneficial mutations to arise; it may, however, alter the spectrum and order of the routes taken. The MDS42 background compounds this: lacking insertion sequences (Pósfai et al. 2006), it excludes IS-mediated promoter remodelling, a common route to efflux upregulation in wild-type *E. coli* (Vandecraen et al. 2017; Nicoloff & Andersson 2013), so the point substitutions and repeat expansions observed upstream of *cmr* may be the accessible alternative in this genome rather than the route that would be taken elsewhere. The landscape was reconstructed in a two-dimensional projection of a three-drug space, a reduction consistent with the low effective dimensionality of resistance phenotypes reported previously (Maeda et al. 2020). It is also an average over heterogeneous lineages, so the region may represent a bottleneck shared across backgrounds or a superposition of lineage-specific constraints, and the sparsely sampled region beyond the ridge is left uninterpreted. Our control comparisons were performed for single-axis targets, and the functional assay was performed in the unevolved ancestor rather than in the evolved multidrug-resistant backgrounds in which the variant naturally acts, where subtle epistatic or energetic costs cannot be excluded. Finally, the expression evidence comes from an upstream variant other than the one we reconstructed, although that variant was itself among those recovered in our evolved lineages.

The particular barrier and genetic route described here are probably specific to this drug set, genetic background and culture condition, but the approach itself is not. Any evolving system in which a phenotype can be quantified, and in which the environment can be adjusted in response to that phenotype, can be examined in the same way: by imposing and recording a known directional perturbation, then estimating local accessibility from the difference between the imposed change and the movement that followed. Used in this way, control becomes an instrument of measurement as well as a means of directing evolution. Beyond antibiotic resistance, the same approach can be applied whenever the question is not only which states evolution can reach, but also where, during adaptation, a population is free to move and where it is not.

## Methods

### Bacterial strains and growth media

All evolution experiments were performed using the insertion-sequence-free, reduced-genome strain *E. coli* K-12 MDS42 (Pósfai et al. 2006) carrying a Δ*mutS* deletion as the ancestral strain. The insertion-sequence-free background was used because the precise position of insertion-sequence transpositions is difficult to determine with short-read sequencing; removing this class of event allows resistance acquisition to be matched to fixed point mutations unambiguously, at the cost of excluding transposition-mediated routes that contribute to resistance evolution in wild-type strains. This Δ*mutS* background was chosen for its high mutation rate to increase mutational supply during laboratory evolution. For the resistance assay shown in Fig. 5a, we additionally used the parental MDS42 strain and an engineered derivative of MDS42 carrying the 746,368 A>G upstream *cmr* mutation, allowing resistance to be compared across parental MDS42, MDS42 Δ*mutS* and MDS42 carrying the upstream *cmr* variant. Unless stated otherwise, cultures were grown at 34 °C in M9 minimal medium supplemented with 15 µg mL^−1^ erythromycin to prevent contamination, a concentration that does not measurably inhibit growth of the ancestral strain, with orbital shaking at 300 rpm.

For routine propagation and revival from frozen stocks, cells were inoculated into liquid cultures and grown overnight before use in evolution experiments or resistance assays. Frozen stocks were prepared by mixing cultures 3:1 with 60% (v/v) glycerol and stored at −80 °C.

### Antibiotics and resistance space definition

To define a three-dimensional resistance space, we used chloramphenicol (CP; Wako 036-10571), norfloxacin (NFLX; Wako 142-06391) and kanamycin sulphate (KM; Wako 117-00341). Antibiotics were prepared as sterile stock solutions in ethanol (CP), acetic acid (NFLX) and M9 medium (KM), respectively, and stored at −20 °C. Working solutions were freshly diluted into prewarmed growth medium immediately before use. Drug-containing medium was prepared at 2,048 µg mL^−1^ (CP), 64 µg mL^−1^ (NFLX) and 2,048 µg mL^−1^ (KM) and held in the reservoirs of the liquid-handling system; these concentrations set the upper bound on the concentrations attainable in culture (see “Adaptive dosing rule and constraints”).

For each antibiotic, we first determined an operational minimum inhibitory concentration (MIC) for the ancestral strain by measuring growth across a series of twofold dilution steps in 96-well plates; the same criterion was subsequently applied to every evolving lineage at each passage. In this study, the operational MIC was defined as the highest concentration at which cultures achieved at least a 100-fold increase in cell density over 24 h at 34 °C, corresponding to the most inhibitory condition that still supported robust growth and was therefore used for passaging. Note that this operational MIC is slightly lower than the classical MIC defined as the lowest concentration with no detectable growth; because it is the highest concentration that still permits growth, it is referred to throughout this study as the sub-MIC. Resistance along each axis was quantified as the log_2_-transformed sub-MIC (hereafter “log_2_ sub-MIC”) or as the log_2_-transformed IC_50_ value (“log_2_ IC_50_”), so that a one-unit difference corresponds to a twofold change in resistance. The resulting three-dimensional space (CP, NFLX, KM) forms the coordinate system in which evolutionary trajectories were steered and analysed.

### Automated culture system

Experimental evolution was conducted using an automated serial-passage platform previously described (Horinouchi et al. 2014). The system consists of a Biomek NXP automated liquid-handling workstation (Beckman Coulter, USA), a FilterMax F3 microplate reader (Molecular Devices, USA; used for all OD_620_ measurements reported here), and an STX44 automated incubator together with an LPX220 plate hotel (Liconic, Liechtenstein). Independent lineages were propagated in microplate culture with daily passages, and antibiotic concentrations were updated in a closed loop based on measured resistance, as described below.

### Control schedules

We implemented seven control schedules that differed in the temporal sequence of resistance targets in the three-drug resistance space (Fig. 1b). In the CP→ALL, NFLX→ALL and KM→ALL schedules, lineages were first steered for 40 days toward high resistance to one focal drug (8 log_2_ µg mL^−1^ for CP and KM, 3 log_2_ µg mL^−1^ for NFLX) while setting the target for the non-focal drugs to their ancestral sub-MICs (CP, NFLX and KM: 1, −5 and 3 log_2_ µg mL^−1^, respectively), and were then steered for 30 days toward the triple-resistance target. In the CP+NFLX→ALL, CP+KM→ALL and NFLX+KM→ALL schedules, lineages were first steered for 40 days toward high resistance to two focal drugs while keeping the remaining drug at its ancestral sub-MIC, and were then steered for 30 days toward the same triple-resistance target. In the ALL→ALL schedule, lineages were steered toward the triple-resistance target throughout the 70-day evolution experiment.

### Daily passage and plate layout

Each day, a small aliquot from each evolving culture was transferred into fresh medium in a 96-well microtiter plate (200 µL per well; initial OD_620_ = 0.0001) and arrayed across wells containing pre-programmed combinations of CP, NFLX and KM concentrations generated by the adaptive dosing algorithm. For each antibiotic, a six-point concentration series was constructed such that the fourth well matched the concentration selected at the previous passage; the remaining wells were filled with drug at concentrations defined by the dilution factor **D** = (*D*_Cℓ_, *D*_NFLX_, *D*_KM_) for each drug. Operationally, for each drug *j*, wells 1–6 corresponded to relative multipliers *D*_j_^+3^, *D*_j_^+2^, *D*_j_^+1^, 1, *D*_j_^−1^, *D*_j_^−2^. Two additional rescue conditions were included at one-half and one-quarter of the fourth-well concentration (wells 7 and 8). The three single-drug series were then overlaid well by well: in well *k* (*k* = 1, ⋯, 8), the concentrations of CP, NFLX and KM were taken from the *k*-th concentration of the corresponding single-drug series. Cells from a given lineage were inoculated into all eight wells of the composite gradient, so that each lineage experienced a full gradient of combined selective pressures around its current operating point. An additional drug-free backup well was included for recovery but was not considered part of the composite multidrug gradient. We refer to the plate carrying these eight combination wells as the combination plate, to distinguish it from the single-drug assay plates described below.

In parallel with this combination plate, aliquots of the same cultures were inoculated into separate single-drug twofold dilution series (12 wells per drug). These assays were used to update the resistance vector **R** for the control algorithm and were later used for IC_50_ estimation.

Plates were incubated for 24 h at 34 °C with orbital shaking (300 rpm), after which OD_620_ was recorded. The propagation decision and resistance update rule are described below.

### Adaptive dosing rule and constraints

To guide evolutionary trajectories toward the desired resistance targets, the per-drug dilution factors were updated by a feedback rule derived from the difference between a target resistance vector **T** = (*T*_CP_, *T*_NFLX_, *T*_KM_) and the current resistance estimate **R** = (*R*_CP_, *R*_NFLX_, *R*_KM_). Here, **R** was computed as the log_2_ sub-MIC from parallel single-drug twofold-dilution assays and was kept conceptually distinct from the operational fourth-well concentrations used to assemble the combination plate. Specifically,

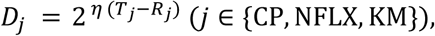

with a scaling factor *η* = 1⁄max {1, ‖**T** − **R**‖} to avoid overly large steps when ‖**T** − **R**‖ > 1. ‖**T** − **R**‖ was at least 1 on 3,638 of the 3,640 lineage-passages, so the vector of log_2_ dilution factors was of unit length: the update had a fixed magnitude and pointed toward the target irrespective of how far the target still lay. On the remaining two passages the measured resistance coincided exactly with the target, so the update was zero. Under this normalisation, *D*_j_ can be greater than or less than 1 and was constrained to the range [2^−1^, 2^+1^], so this bound was never active.

The absolute concentrations were nevertheless bounded from above by the dilution scheme itself. Drug-containing medium was supplied from reservoirs at 2,048 µg mL^−1^ (CP), 64 µg mL^−1^ (NFLX) and 2,048 µg mL^−1^ (KM), and the smallest dilution of this medium that could be prepared in a well was 0.267-fold (4/15). The highest concentration attainable in any well was therefore 546.1 µg mL^−1^ (9.09 log_2_ µg mL^−1^) for CP, 17.07 µg mL^−1^ (4.09 log_2_ µg mL^−1^) for NFLX and 546.1 µg mL^−1^ (9.09 log_2_ µg mL^−1^) for KM; these ceilings are marked by the black dashed lines in Supplementary Fig. 1. Each lies 1.09 log_2_ units above the corresponding high-resistance target level (8, 3 and 8 log_2_ µg mL^−1^ for CP, NFLX and KM), so every assigned target lay within the attainable range, and when the dosing rule called for a concentration above the ceiling, the ceiling concentration was used instead. As a lineage’s resistance approached the target along a given axis, the corresponding component of the update approached zero, so that the unit-length step was carried by the remaining axes.

### Resistance estimation and propagation decision

At each daily passage, single-drug twofold dilution assays were used both for real-time feedback control and for subsequent quantitative analysis of resistance trajectories. The log_2_ sub-MIC estimated from these assays was used to update the adaptive dosing rule for the next passage, whereas the full dose–response profiles from the same assays were later fitted to estimate IC_50_ values for CP, NFLX and KM. Each assay consisted of a 12-point twofold concentration series for each drug, spanning 0.125–256 µg mL^−1^ for CP, 0.0039–8 µg mL^−1^ for NFLX and 0.125–256 µg mL^−1^ for KM. For each lineage, time point and drug, growth across this concentration series was measured after 24 h incubation at 34 °C with shaking.

For the combination plate, each lineage was inoculated into wells containing the composite CP–NFLX–KM gradient generated from the current fourth-well concentrations and the drug-specific dilution factors. After 24 h culture, wells with OD_620_ > 0.01, corresponding to the 100-fold increase used to define the sub-MIC, were considered to show detectable growth. Among the six dynamically updated gradient wells exceeding this threshold, the highest-priority viable well, defined as the lowest-index well in the composite gradient, was selected for propagation, and the drug concentrations in this selected well were used as the fourth-well concentrations for the next passage.

If none of the six dynamically updated gradient wells showed detectable growth, propagation was attempted from the two lower-concentration rescue wells, in which all three drugs were reduced to one-half or one-quarter of the fourth-well concentrations. The higher-concentration rescue well was prioritised. If these rescue wells also failed to show detectable growth, propagation was performed from an additional drug-free well, and the fourth-well concentrations for the next passage were set to one-eighth of those of the current passage, that is, half the concentration of the lower rescue well. In rare cases where no growth was detected in any of these wells, the passage was repeated using the drug-free well inoculated from the residual culture of the previous passage, which had been stored at 4 °C, and the repeated passage used fourth-well concentrations one-eighth of those of the preceding passage.

### Target-distance analysis

Target approach was quantified in the three-dimensional log_2_ sub-MIC resistance space used for real-time feedback control. For each lineage and time point, the resistance state was represented as **R** = (*R*_CP_, *R*_NFLX_, *R*_KM_), and its distance to a target state **T** = (*T*_CP_, *T*_NFLX_, *T*_KM_) was computed as the Euclidean distance ‖**R** − **T**‖.

For first-phase target analyses, day-40 resistance states were compared with the assigned first-phase target and with the other candidate target states. This analysis was used to assess whether each target schedule drove lineages preferentially toward its assigned direction in resistance space. For the triple-resistance analysis, day-70 resistance states were compared with the common ALL target, and the distribution of distances across lineages was used to quantify incomplete convergence to the target region. To test whether day-70 endpoint positions depended on the first-phase schedule, we applied permutational multivariate analysis of variance (PERMANOVA; Anderson 2001) to Euclidean distances among day-70 states, in both the three-dimensional log_2_ IC_50_ space and the PCA plane, with 9,999 permutations, and tested homogeneity of multivariate dispersion using PERMDISP (Anderson 2006). Both tests were performed with the adonis2 and betadisper functions of the R package vegan (Oksanen et al. 2022). Differences among schedules in the day-70 distance to the ALL target were tested by the Kruskal–Wallis test. To quantify preferential approach across the whole design, for each of the 28 first-phase lineages we ranked the seven candidate targets by Euclidean distance from the day-40 state (rank 1 = nearest) and used the mean rank of the assigned target across lineages as the test statistic. An exact null distribution was obtained by enumerating all 5,040 bijective assignments of the seven schedules to the seven targets, yielding an exact *P* value; per-schedule significance was assessed by an analogous exact test with Benjamini–Hochberg correction across the seven schedules.

For control experiments comparing single-drug evolution, dynamic three-drug control and fixed-direction three-drug control, distances from day-40 resistance states to the assigned single-axis target were computed in the same log_2_ sub-MIC space (*n* = 12 lineages per scheme, pooling the three single-axis target conditions). The effect of control scheme was tested by a two-way analysis of variance with control scheme and target condition as factors, and confirmed by permutation of the scheme labels within each target condition (9,999 permutations); pairwise contrasts between schemes were Benjamini–Hochberg corrected. For the pooled display in Supplementary Fig. 2b, each distance was adjusted for its target condition as *d̅_ik_* = *d_i_*_k_ − (*d̅*_k_ − *d̅*), where *d̅*_k_ is the mean distance for target condition *k* across all three schemes and *d̅* is the grand mean; this removes the target main effect without altering the differences between schemes, and was used for display only. In the fixed-direction control, only the direction of the concentration update was fixed at its initial value, while its magnitude continued to follow growth, isolating the contribution of directional updating.

### Resistance assays and IC_50_ estimation

For quantitative analysis of resistance trajectories, the same single-drug twofold dilution assay data used for estimating log_2_ sub-MIC values were also used to estimate IC_50_ values for CP, NFLX and KM. The dose–response profiles from these assays (12-point twofold series, 24 h at 34 °C with shaking) were fitted as follows. The dose–response curve was fitted on the log_2_ concentration scale using the following sigmoidal model:

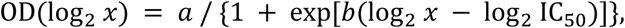

where *x* is the drug concentration, *a* is the fitted growth amplitude, *b* is the slope parameter, and log_2_ IC_50_ is the fitted log_2_ concentration at which growth is reduced by half relative to the fitted amplitude. Fitting was performed independently for each lineage, time point and drug. Fits were visually inspected, and no fitting failures requiring exclusion were observed. The resulting log_2_ IC_50_ values were used for visualising trajectories in three-dimensional log_2_ IC_50_ space and for downstream PCA-based landscape reconstruction.

### PCA and reconstruction of the accessibility landscape

For PCA and landscape reconstruction, log_2_ IC_50_ values were normalised separately for each drug by subtracting the mean and dividing by the standard deviation across all lineage–time data points. The normalised three-dimensional resistance states were then projected onto the first two principal components. PC1 primarily captured variation in CP and NFLX resistance, whereas PC2 primarily captured variation in KM resistance. The first two principal components explained 58% and 32% of the variance, respectively.

We analysed N = 3,588 lineage-day intervals, corresponding to 52 lineages over 69 daily transitions. For each lineage *_ℓ_* and day *t*, the positions before and after one 24 h culture step in the PCA-transformed log_2_ IC_50_ space were denoted as **x***_ℓ_*_,*t*_ and **x***_ℓ_*_,*t*+1_, respectively, and the realised displacement was defined as

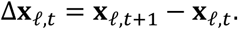

For each lineage-day interval, the target-directed dilution update produced drug-specific dilution factors **D** = (*D*_CP_, *D*_NFLX_, *D*_KM_). The vector of log_2_ dilution factors, (log_2_ *D*_CP_, log_2_ *D*_NFLX_, log_2_ *D*_KM_), was treated as a nominal controller-defined input describing the direction and relative size of the concentration update in the original CP–NFLX–KM coordinate system. To compare this controller-defined input with the observed displacement in the PCA plane, this vector was scaled by the same drug-wise standard deviations used to normalise log_2_ IC_50_ values and then projected using the same PCA loading matrix. The resulting two-dimensional vector was denoted as **q***_ℓ_*_,*t*_. Thus, **q***_ℓ_*_,*t*_ encoded both the direction and relative magnitude of the imposed control update in the PCA-transformed log_2_ IC_50_ coordinate system. To match the overall scale of the control vectors to that of the observed displacements, we multiplied **q***_ℓ_*_,*t*_ by a global scaling factor *λ*, defined as

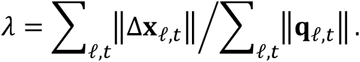

In this dataset, *λ* = 0.549. The residual vector, representing the component of phenotypic movement not explained by the applied control vector, was then defined as

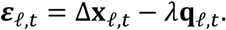

This formulation treats the log_2_ fold change in drug concentration as a nominal controller-defined input, not as a calibrated measure of selection strength. Thus, **q***_ℓ_*_,*t*_ defines a reference direction generated by the dosing rule: log_2_ *D*_j_ > 0 corresponds to a concentration update toward higher resistance along drug *j*, log_2_ *D*_j_ = 0 corresponds to no directional update, and log_2_ *D*_j_ < 0 corresponds to relaxation toward lower concentrations. *λ***q***_ℓ_*_,*t*_ therefore represents the globally scaled reference component, whereas drug-specific or state-dependent deviations from the nominal input are retained in the residual vector ***ε****_ℓ_*_,*t*_ and contribute to the inferred landscape only when they show reproducible spatial structure; the analysis does not assume that identical fold changes in concentration exert identical phenotypic effects or selection strengths across drugs. The global gain was estimated by moment matching as the ratio of summed observed displacement lengths to summed control-vector lengths (0.549). The sensitivity of the residual field to this value was assessed by repeating the reconstruction at *λ* = 0, *λ*/2 and 2*λ*; the results, and their consequences for the interpretation of the fitted landscape, are given in Results and Supplementary Fig. 5e.

We interpreted these residual vectors as local deflections in resistance space and fitted a scalar landscape *L*(**x**) whose gradient accounted for them. The landscape was represented as a grid of non-negative height values *L_n_*_,*m*_ on the PCA plane. The grid widths along PC1 and PC2 were determined using the Freedman–Diaconis rule (Freedman & Diaconis 1981) and were set to *b*_1_ = 0.27 and *b*_2_ = 0.21, respectively. The grid was defined so that the forward finite differences used below were evaluable for all occupied cells. For each position **x***_ℓ_*_,*t*_ assigned to grid cell (*n*, *m*), the local gradient was evaluated by forward finite differences:

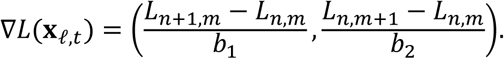

The landscape was fitted by minimising the discrepancy between the landscape-derived direction and the residual vectors:

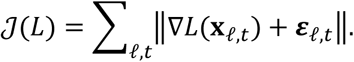

Here, the vector norm was computed as the Euclidean norm. This objective corresponds to fitting *L*(**x**) such that −∇*L*(**x***_ℓ_*_,*t*_) aligns with ***ε****_ℓ_*_,*t*_. Optimisation was performed using scipy.optimize.basinhopping (Virtanen et al. 2020) with 100 iterations and the default L-BFGS-B local minimiser. Initial landscape heights were randomly assigned between 0 and 1, and the optimisation was constrained to keep all grid heights positive. After convergence, landscape heights ranged from 0.02 to 0.87 before the Gaussian smoothing described below. A grid cell was included in the analysis only if at least one independent lineage contributed to it, that is, if the cell contained a pre-culture or post-culture position of that lineage, or lay in the forward target-direction neighbourhood used to evaluate the finite-difference gradient; cells to which no independent lineage contributed were masked and left uninterpreted.

To assess whether the inferred landscape captured reproducible spatial structure rather than overfitting, we performed held-out validation at the lineage level. The 52 lineages were split at random into a training set of 39 and a held-out set of 13. The landscape was reconstructed from the training lineages and then evaluated on the held-out lineages, corresponding to 897 lineage-day intervals. For a two-dimensional vector **v** in the PCA plane, *θ***_v_** denotes its direction, and ***ε***_test_ denotes the residual vectors evaluated on the held-out lineages; the quantity plotted in Fig. 2c is the absolute angular difference |*θ****_ε_*** − *θ*_−∇*L*_|, wrapped to [0, π]. We compared the mean of this quantity with a null distribution obtained from B = 9,999 permutations of the residual vectors among the held-out positions. No permutation was as well aligned as the observation (observed mean 1.26 rad versus null mean 1.49 rad), giving a one-sided *P* < 1/(B + 1) = 10⁻⁴ (z = −7.9); the same held when permutation was restricted within lineages (z = −6.5) or replaced by within-lineage cyclic shifts along the day axis, which preserve temporal correlation (z = −3.9; both *P* < 10⁻⁴).

### Whole-genome sequencing and variant calling

For whole-genome sequencing, samples were taken from the three target schedules with the largest number of replicate lineages: KM→ALL, CP+NFLX→ALL and ALL→ALL, with 12 independent lineages per schedule. The two time points were sampled in different ways. At day 40, genomic DNA was extracted from the evolving population without single-colony isolation. At day 70, each population was streaked for single colonies; four colonies were picked per lineage, their log₂ sub-MIC values were determined for CP, NFLX and KM as in the daily assay, and the colony whose resistance vector lay closest to that of the day-70 population was sequenced. Day-70 genotypes therefore describe one clone chosen to represent the population endpoint, whereas day-40 genotypes describe the population as a whole. The two time points were compared to distinguish mutations that arose during the initial target phase from those that were present or enriched during the subsequent approach to the common triple-resistance target. In addition, four drug-free control lineages propagated for 70 days were sequenced at the same two time points to identify mutations that accumulated reproducibly during serial passage in the absence of drug selection.

Evolved population stocks were revived under the standard culture conditions described above and grown until sufficient cell density was reached for DNA extraction. Cells were harvested by centrifugation, and genomic DNA was extracted using a DNeasy Blood & Tissue Kit (Qiagen). Sequencing libraries were prepared using a Nextera XT kit (Illumina) and sequenced on an Illumina MiSeq platform in a paired-end 2 × 300 bp setting using the MiSeq Reagent Kit v3 (600-cycle). Short reads were mapped to the *E. coli* K-12 MDS42 reference genome (GenBank accession: AP012306.1), and mutations were detected using breseq v0.35.1 (Deatherage & Barrick 2014) in its default consensus mode with bowtie2 v2.1.0 (Langmead & Salzberg 2012) and R v3.2.4. Detected variants were annotated according to their positions relative to coding sequences and intergenic regions. The full list of detected variants is provided in Supplementary Data 1 (sheet “Detected variants”).

To focus on mutations associated with the present drug-selection experiment, background or experiment-wide variants were excluded before downstream analyses. These included five variants common to the laboratory MDS42 ancestral background, 21 variants already present in the Δ*mutS* ancestral strain used for this experiment, and one recurrent *ydgJ*–*blr* intergenic variant that was broadly observed in the present experiment and was also fixed in drug-free control lineages. The excluded background and experiment-wide variants are listed in Supplementary Data 1 (sheet “Excluded variants”). After these exclusions, remaining mutations were grouped by affected gene or intergenic region for downstream analyses.

In addition, the day-40 alignments were re-examined at read level at the *ybjG*–*cmr* locus, because consensus calling reports only variants that have reached majority frequency in the population sample (Supplementary Note 1).

### Barrier definition and crossing classification

After reconstructing the accessibility landscape using all 52 lineages, the fitted landscape was smoothed using a Gaussian kernel with bandwidth ℎ in PCA-coordinate units. Candidate bandwidths were evaluated by four-fold cross-validation over the grid cells on which the landscape is defined, independently of the held-out validation shown in Fig. 2c. The 396 such cells were divided at random into four groups of 99. For each candidate bandwidth, the Gaussian smoothing was fitted from the cells of three groups, and the squared difference between the raw landscape height and the smoothed height was accumulated over the cells of the held-out group. Summed over the four folds and divided by the number of cells in one group (99), this is the cross-validation error plotted in Supplementary Fig. 6a. The bandwidth minimising it, ℎ = 0.32, was used to smooth the final landscape reconstructed from all 52 lineages.

To define the barrier line, we considered a diagonal reference line in the PCA plane running from the upper-left to the lower-right side of the plotted region. From grid cells along this reference line, we scanned in the perpendicular direction toward the upper-right, corresponding to the target-side region in this PCA representation. Along each scan line, the first grid cell at which the smoothed landscape value decreased relative to the preceding cell was marked as a candidate barrier cell, corresponding to a local ridge in *L*. Where a gap remained between two candidate barrier cells that were diagonal neighbours, a single bridging cell was inserted to obtain a continuous boundary; two such cells were required at the gain used for the analyses reported here (Supplementary Fig. 6b). The connected boundary defined by these candidate and bridging cells was used as the barrier line.

The side of this barrier line containing the projected triple-resistance target was defined as the post-barrier region, referred to as the target-side region in Results, whereas the opposite side was defined as the pre-barrier region. For each lineage, the PCA-transformed log_2_ IC_50_ trajectory was overlaid on this fixed barrier definition. Lineages were classified as Crossed if their trajectory entered the post-barrier region at least once during the 70-day evolution experiment; otherwise, they were classified as Stalled. This classification was used for the mutation-enrichment analysis.

To test whether the classification depends on the global gain, the landscape was refitted at *λ*/2 and 2*λ*, smoothed with the same bandwidth (ℎ = 0.32), and the ridge-detection and bridging procedure was attempted unchanged. Because *L* is fitted independently at each gain, its absolute height is not comparable across gains; only the position of the ridge and of the resulting boundary is. At 2*λ* the procedure returned a continuous boundary that coincided cell for cell with the one obtained at *λ*, so every lineage received the same classification and every analysis that rests on it was unchanged. At *λ*/2 the detected candidate cells left gaps wider than the single-cell bridging criterion above, so no boundary separating the sampled region was obtained, and the two sides of a barrier are undefined at that gain. The gaps could be closed by hand, but the position of the resulting boundary, and the classification of several lineages with it, would then depend on where the connecting cells were placed rather than on the fitted landscape; the classification is therefore reported at *λ* and 2*λ* only.

### Mutation enrichment analysis

After exclusion of background and experiment-wide variants, mutations were grouped by affected gene or intergenic region. Loci mutated in at least five of the 36 drug-treated sequenced lineages were included in the enrichment analysis. For each locus, a 2 × 2 contingency table was constructed from the number of Crossed and Stalled lineages with or without a mutation in that locus at day 70. Enrichment in Crossed lineages was tested using Fisher’s exact test. *P* values were corrected for multiple testing using the Benjamini–Hochberg false discovery rate procedure (Benjamini & Hochberg 1995), and loci with FDR < 0.05 were considered significantly enriched. Effect sizes for enriched loci were summarised as Haldane–Anscombe-corrected 95% confidence intervals. To corroborate the association independently of the Crossed/Stalled classification, we also compared the day-70 distance to the triple-resistance target in the PCA plane between lineages with and without an upstream *cmr* variant across all 36 sequenced lineages, using a two-sided Mann–Whitney U test with Cliff’s δ (Cliff 1993) as the effect size (Supplementary Fig. 7). As a negative control, the four drug-free control lineages propagated for 70 days were examined for upstream *cmr* variants; none was detected.

### Analysis of upstream *cmr* variants and expression datasets

To analyse mutations upstream of *cmr*, variants detected in the *ybjG*–*cmr* intergenic region were mapped onto the *E. coli* K-12 MDS42 genome coordinate system. Their positions were compared with the coding directions of *ybjG* and *cmr* (*mdfA*) and with annotated or predicted regulatory features in this region. Variants were classified by type, including single-nucleotide substitutions, a short repeat expansion and closely spaced substitutions.

To examine whether upstream *cmr* variants were associated with altered *cmr* expression, we re-analysed a previously published microarray dataset from laboratory evolution experiments under chloramphenicol and puromycin selection (Maeda et al. 2020). Expression values were taken from the normalised microarray data provided in the original study. In that dataset, four independently evolved lineages were available for each drug condition. For each of the chloramphenicol and puromycin experiments, one lineage carried the upstream *cmr* variant 746,309 (TAATGTAA)_1→2_, whereas three lineages lacked this variant. We compared *cmr* expression between the variant-carrying lineage and the three non-carrier lineages within each drug condition, and visualised the values in Supplementary Fig. 8b. Because each drug condition contained only one variant-carrying lineage, this comparison was treated as a descriptive re-analysis of the published expression dataset.

### Construction of upstream *cmr* mutant strains and resistance phenotyping

To test the phenotypic effect of the recurrent upstream *cmr* mutation, the 746,368 A>G variant was introduced into the unevolved *E. coli* K-12 MDS42 background using multiplex automated genome engineering (MAGE; Wang et al. 2009), following the procedure described in Maeda et al. (2020). The oligonucleotide used for genome editing is listed in Supplementary Table 1. The resulting mutant genotype was confirmed by Sanger sequencing.

Resistance phenotypes of the parental MDS42 strain and the engineered upstream *cmr* mutant were quantified using the same single-drug twofold dilution assay described above. Briefly, growth was measured across 12-point twofold concentration series for CP, NFLX and KM after 24 h incubation at 34 °C with shaking, and IC_50_ values were estimated by fitting the dose–response curves on the log_2_ concentration scale. The MDS42 Δ*mutS* strain was included as an additional control; its IC_50_ values differed from parental MDS42 by at most 0.17 log_2_ units on any axis, less than one twofold dilution step, although this difference was detectable at the precision of the assay (*n* = 8, *P* < 0.001). The minimum difference detectable on the NFLX and KM axes was estimated from the replicate standard deviation of the log_2_ IC_50_ values for a two-sample comparison (*n* = 8 per strain) at 80% power (two-sided, *α* = 0.05).

Growth rates of the parental MDS42 strain and the engineered upstream *cmr* mutant were measured in medium without CP, NFLX or KM under the same culture conditions used for the resistance assay. Overnight cultures were diluted into fresh prewarmed M9 medium and inoculated into 96-well plates. Cultures were incubated at 34 °C with shaking in an Infinite 200 PRO microplate reader (Tecan, Switzerland), and optical density at 600 nm (OD_600_) was recorded every 15 min. Growth rates were estimated from the exponential phase of each growth curve. Values were summarised across biological replicates and used for Supplementary Fig. 8a.

For comparison with other resistance-associated mutations, IC_50_ effects of previously reconstructed representative mutant strains were taken from the published dataset of Maeda et al. (2020), generated using the same MAGE-based reconstruction and resistance-phenotyping framework. For each mutant, resistance effects were expressed as fold changes in IC_50_ relative to the corresponding parental strain. These values were used to compare the CP, NFLX and KM resistance effects of the upstream *cmr* mutation with those of other representative resistance-associated mutations. All 64 reconstructed mutants from the Maeda et al. (2020) panel were used, with the 14 mutants in gene regions overlapping recurrently mutated loci in our lineages highlighted. To quantify how distinctive the upstream *cmr* phenotype was, we counted panel mutants whose CP gain matched or exceeded that of the upstream *cmr* variant (≥ 1.55 log_2_ units) without a decrease along the NFLX or KM axes, and computed an empirical *P* value as (*k* + 1)/(64 + 1).

## Statistical analysis

Statistical procedures specific to each analysis are described in the corresponding Methods sections. Unless otherwise stated, tests were two-sided, and summary values are reported as mean ± standard deviation.

## Data availability

Sequencing data generated in this study have been deposited in the DDBJ Sequence Read Archive under BioProject accession PRJDB42845 (BioSample accessions SAMD01947744–SAMD01947823; run accessions DRR1079331–DRR1079410). The processed datasets generated and analysed in this study — the daily sub-MIC, mixed-drug MIC, IC_50_ and dilution trajectories of every lineage, the target trajectories, the drug schedules, the variant table and its gene annotation, and the IC_50_ and growth rate of the reconstructed strains — are available in the Zenodo repository at https://doi.org/10.5281/zenodo.22074247. The resistance phenotypes of the reconstructed mutants re-analysed in Fig. 5b, and the gene expression and mutation data of evolved strains re-analysed in Supplementary Fig. 8b, are the supplementary data files of Maeda et al. (2020) and are available with that paper (https://doi.org/10.1038/s41467-020-19713-w). Source data are provided with this paper.

## Code availability

All code used to produce the figures and the statistics reported here is available in the Zenodo repository at https://doi.org/10.5281/zenodo.22074247 (Shibai et al. 2026). Each figure panel is produced by a pair of scripts: one that turns the measurements into the Source Data table of that panel, and one that draws the panel from that table alone. The code is released under the MIT License and the accompanying data under the Creative Commons Attribution 4.0 International License.

## Acknowledgements

We thank Shumpei Sato and Naomi Yokoi for helpful discussions and technical support. This work was supported by the JST ACT-X program (JPMJAX20B6 to A.S.), the JST PRESTO program (JPMJPR25T4 to A.S.) and the JST ERATO program (JPMJER1902 to C.F.). It was also supported by JSPS KAKENHI (21K15077 to A.S.; 17H03622, 22K21344, 23H02471, 26H01831 to C.F.), the Uehara Memorial Foundation and the Mitsubishi Foundation.

## Author contributions

A.S. and C.F. conceived the study. A.S. designed and performed the experiments. A.S. and H.K. performed genome sequencing and related analyses. A.S. analysed the data. A.S. and C.F. interpreted the results. A.S. wrote the initial manuscript draft. All authors reviewed and edited the manuscript.

