## Supplementary Information for "Closed-loop steering locates an evolutionary barrier to multidrug resistance in *Escherichia coli*"

[*]**Corresponding author**

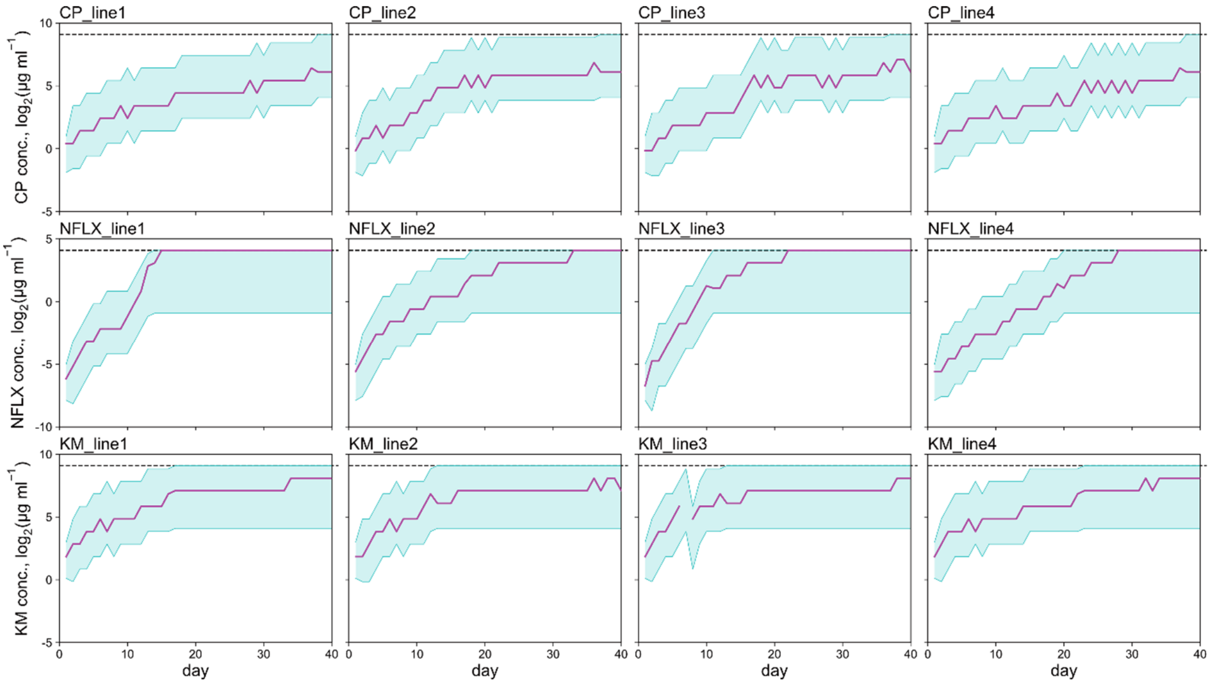


**Supplementary Fig. 1 | Single-drug closed-loop evolution under CP, NFLX and KM.**

Trajectories of $\log_2$ sub-MIC against time (horizontal axis, days) for four independent MDS42 Δ*mutS* lineages per drug, evolved for 40 days under chloramphenicol (CP, top row), norfloxacin (NFLX, middle row) or kanamycin (KM, bottom row). In each lineage, the sub-MIC (magenta lines) increased over time, causing the six-step dilution range offered at each passage (cyan shading; the six dynamically updated gradient wells, Fig. 1c) to shift upward. The black dashed line in each panel marks the maximum concentration attainable in culture, set by the reservoir concentration and the smallest available dilution (546.1, 17.07 and 546.1 µg $mL^{-1}$ for CP, NFLX and KM, respectively; Methods). For NFLX, sub-MIC values reached this limit between about days 12 and 27 and subsequently plateaued near it. In the third KM lineage, the break in the trajectory at day 7 reflects missing data rather than a true decrease in resistance. These single-drug experiments were performed separately from the 52 lineages of the main experiment. Source data are provided with this paper.

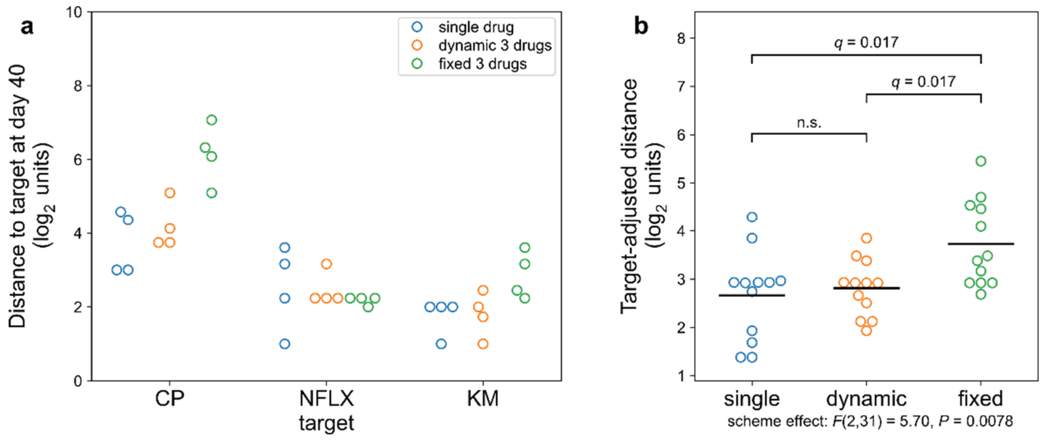


**Supplementary Fig. 2 | Comparison of single-drug, dynamic three-drug and fixed three-drug control schemes.**

For each single-axis target condition (CP, NFLX or KM), day-40 resistance states were obtained under three schemes: single-drug evolution; dynamic three-drug control, in which the concentration gradient was updated each passage from the difference between the current state and the target; and fixed-direction three-drug control, in which the direction of the update was fixed at its initial value while its magnitude continued to follow growth. **a**, Euclidean distance between each day-40 state and its assigned single-axis target in $\log_2$ sub-MIC space, shown separately for each target condition; each point represents one lineage (four per scheme and target) and colours denote the control scheme. **b**, The same distances after adjustment for the target condition (target-adjusted distance; Methods), pooled across the three targets ($n = 12$ lineages per scheme); colours as in **a**, and horizontal bars show the group mean. The effect of scheme was tested by two-way analysis of variance blocking on target (scheme main effect $F(2,31) = 5.70$, $P = 0.0078$) and confirmed by within-target permutation ($P = 0.0083$); pairwise contrasts were Benjamini–Hochberg corrected and are reported as q, the adjusted P value. Fixed-direction control reached significantly farther from the target than single-drug evolution (+1.07 $\log_2$ units, $q = 0.017$) or dynamic control (+0.92 $\log_2$ units, $q = 0.017$), whereas dynamic control and single-drug evolution did not differ (+0.15 $\log_2$ units, $q = 0.61$). Source data are provided with this paper.

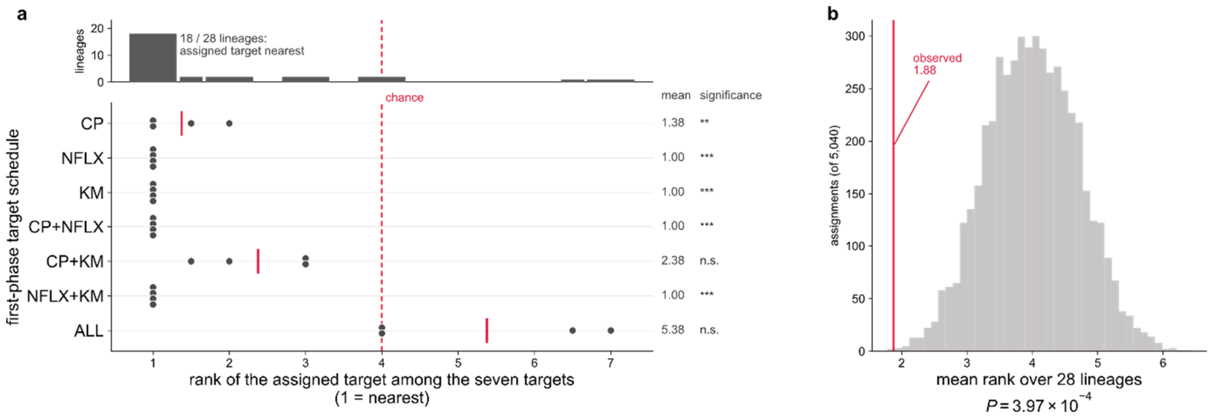


**Supplementary Fig. 3 | Day-40 states are closest to their assigned first-phase target.**

For each of the 28 first-phase lineages, comprising four lineages from each of the seven schedules so that the schedules are equally represented, the seven candidate targets were ranked by Euclidean distance from the day-40 state in three-dimensional $\log_2$ sub-MIC space (rank 1 = nearest). **a**, Rank of the assigned ("own") target for each lineage, grouped by schedule; the histogram (top) shows that the own target was the nearest for 18 of 28 lineages, and the dashed line marks the chance expectation (rank 4). Red bars mark per-schedule means, and per-schedule significance (exact test, Benjamini–Hochberg corrected across the seven schedules) is indicated at right as adjusted P values: $^{**}q < 0.01$, $^{***}q < 0.001$; n.s., not significant. The own target was significantly closer than the non-self targets in five of the seven schedules. The two exceptions, CP+KM and ALL, are the only two targets requiring simultaneous increases along both the CP and KM axes and are consistent with the collateral sensitivity between CP and KM. **b**, Exact null distribution of the mean own-target rank over all 5,040 schedule-to-target assignments; the observed mean (1.88) lies far below the chance expectation ($P = 3.97 \times 10^{-4}$). Source data are provided with this paper.

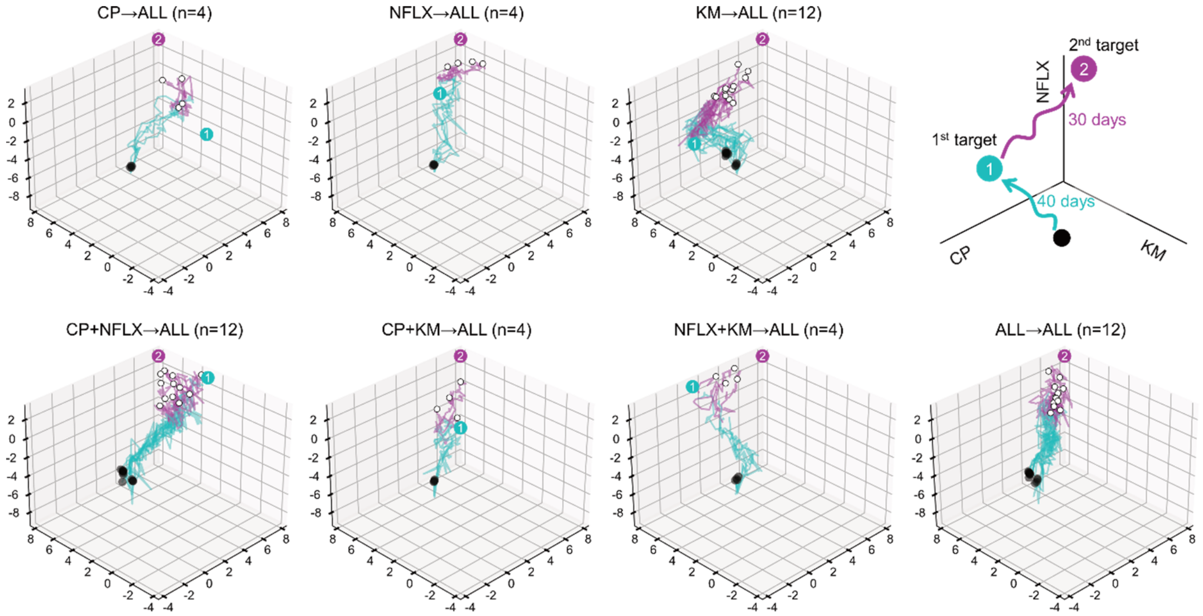


**Supplementary Fig. 4 | Evolutionary trajectories in three-dimensional $\log_2$ $IC_{50}$ space.**

The same evolutionary trajectories shown in Fig. 1b were replotted using $\log_2$ $IC_{50}$ values estimated from dose–response assays; n is the number of independent lineages per schedule. Cyan indicates the first 40 days toward the assigned first-phase targets (label ①), and magenta indicates the subsequent 30 days toward the common triple-resistance target (ALL, label ②). In ALL→ALL, the two targets coincide; thus, only the second is shown. Black spheres indicate the ancestral starting point (day 1). White spheres indicate day-70 endpoints. The schematic at the top right shows the two-phase design and the orientation of the CP, NFLX and KM axes, which is the same in every subpanel and as in Fig. 1b; axes show $\log_2$ $IC_{50}$ (μg $mL^{-1}$). Source data are provided with this paper.

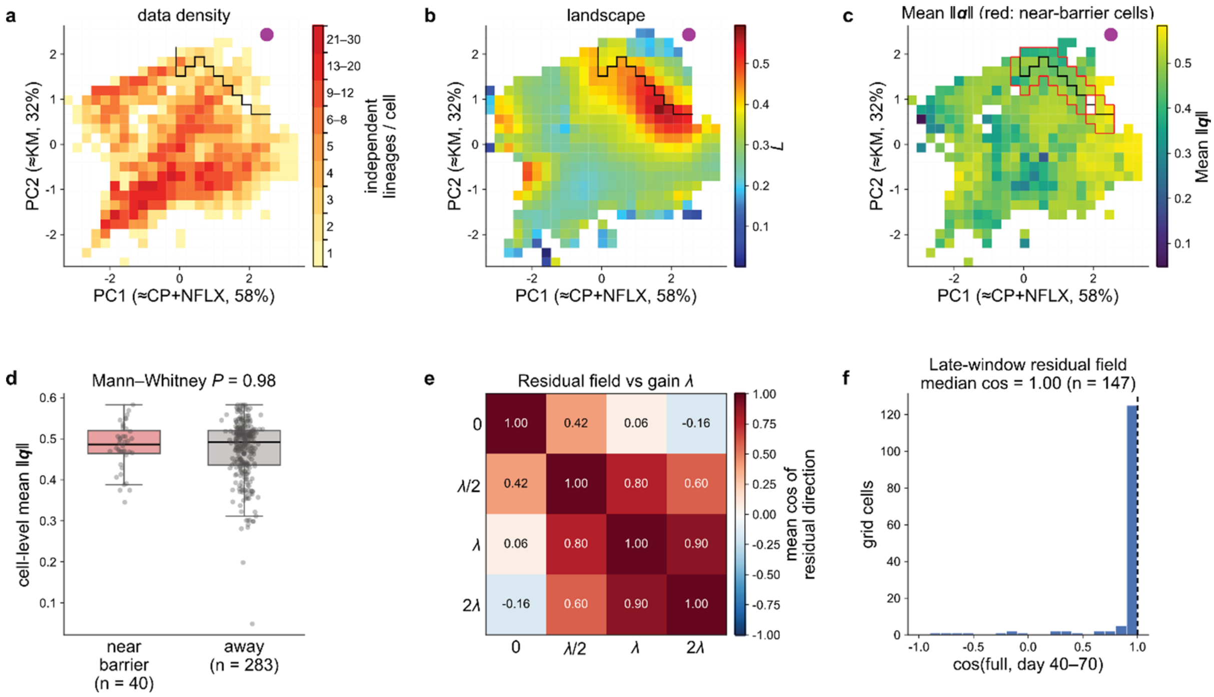


**Supplementary Fig. 5 | Robustness of the reconstructed accessibility landscape.**

Panels a–c are shown in the PCA plane of normalised $\log_2$ $IC_{50}$ values (PC1 ≃ CP+NFLX, 58%; PC2 ≃ KM, 32%), in which the black line is the barrier defined in Methods and the magenta circle marks the projected triple-resistance target. **a**, Number of independent lineages contributing to each grid cell. The barrier and ridge lie within the densely sampled region, whereas the target-side corner near the circle is sparsely sampled, which is why the apparent decrease of $L$ beyond the ridge is not interpreted. **b**, Reconstructed accessibility landscape $L$ (as in Fig. 2d); the ridge in $L$ coincides with the barrier line. **c**, Mean magnitude of the projected control vector, $\|\mathbf{q}\|$, per grid cell; near-barrier cells are outlined in red. **d**, Cell-level mean $\|\mathbf{q}\|$ does not differ between near-barrier cells (pink, n = 40) and the remaining cells (grey, n = 283) (two-sided Mann–Whitney U test, $P = 0.98$), so the barrier does not coincide with a region of systematically weaker or stronger control input. Boxes show the median and interquartile range with whiskers, and points show individual grid cells. **e**, Sensitivity of the residual field to the global gain $\lambda$. For each $\lambda$ (0, $\lambda/2$, $\lambda = 0.549$ and $2\lambda$), the per-cell mean residual direction was computed; the matrix shows the mean cosine between the residual-direction fields of each pair of values, weighted by the number of lineage-day intervals contributing to each cell. Over the fourfold range from $\lambda/2$ to $2\lambda$ the fields agree closely (mean cosine 0.60 to 0.90 across the three pairs), whereas the field at $\lambda = 0$, in which the control input is discarded, is unrelated to the fields at $\lambda$

and 2$\lambda$ (0.06 and −0.16, respectively), although its cosine against the field at $\lambda/2$ is 0.42. The residual field is therefore insensitive to the precise value of the gain once the control input is retained, but not to omitting it; the robustness of the ridge and of the barrier drawn from it is reported separately in Methods. **f**, The residual-direction field reconstructed from the day-40 to day-70 interval alone agrees with that from the full time series (median per-cell cosine ≈ 1.0; n = 147 cells), indicating that the reduced-accessibility structure is not an artefact of the earlier, first-phase portion of the trajectories. Source data are provided with this paper.

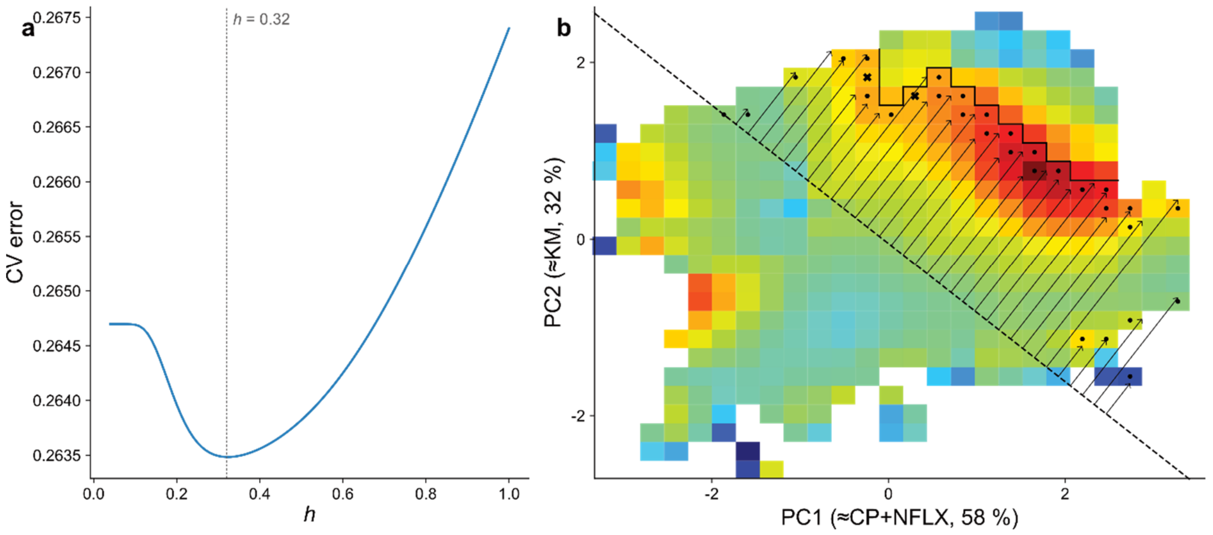


**Supplementary Fig. 6 | Definition of the reduced-accessibility region (barrier) on the reconstructed accessibility landscape.**

**a**, Selection of the Gaussian smoothing bandwidth $h$. The final landscape reconstructed from all 52 lineages was smoothed using a Gaussian kernel in PCA-coordinate units. The cross-validation error (CV error) is plotted against the candidate bandwidth. The bandwidth was selected by four-fold cross-validation over the grid cells on which the landscape is defined: the cells were divided at random into four groups, the smoothing was fitted from three groups, and the squared error was accumulated on the held-out group (Methods). This is independent of the held-out validation shown in Fig. 2c. The bandwidth minimising the CV error was $h = 0.32$. **b,** Procedure used to define the barrier line, shown on the smoothed landscape; colours show the smoothed landscape $L$ as in Fig. 2d, with warmer colours denoting higher $L$. The dashed line indicates the diagonal reference line used for ridge detection. From grid cells along this line, scans were performed in the perpendicular direction toward the post-barrier region, as indicated by the arrows. Dots indicate automatically detected candidate barrier cells where the smoothed landscape value first decreased along each scan line. Crosses indicate the two bridging cells added to obtain a continuous boundary. The solid black line indicates the resulting barrier used to define the post-barrier region, and is the same barrier shown in Fig. 2d. Source data are provided with this paper.

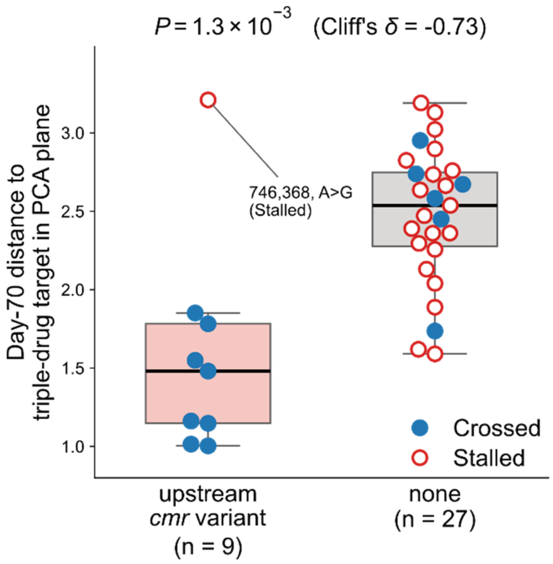


**Supplementary Fig. 7 | Upstream *cmr* variants are associated with target proximity independently of the barrier classification.**

Day-70 distance to the triple-resistance target in the PCA plane for all 36 sequenced lineages, grouped by whether they carry an upstream *cmr* variant (n = 9) or not (n = 27). Filled blue symbols denote lineages classified as Crossed and open red symbols denote Stalled lineages; boxes show the median and interquartile range with whiskers. Lineages carrying an upstream *cmr* variant ended closer to the target regardless of their Crossed/Stalled classification (median distance to the target 1.5 versus 2.5 units; two-sided Mann–Whitney U test, $P = 1.3 \times 10^{-3}$; Cliff's $\delta = -0.73$), showing that the association in Fig. 4c does not depend on the barrier definition. The one carrier of the recurrent 746,368 A>G variant that remained far from the target (indicated by the annotation) was classified as Stalled, consistent with this variant being neither necessary nor sufficient for barrier crossing. Source data are provided with this paper.

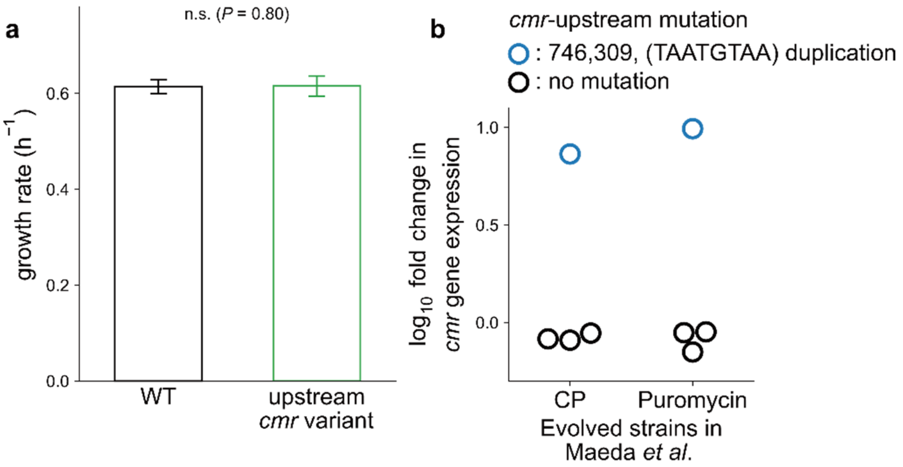


**Supplementary Fig. 8 | Growth and expression analyses supporting the functional interpretation of upstream *cmr* variants.**

**a,** Growth rates of parental MDS42 (WT) and the engineered MDS42 strain carrying the 746,368 A>G upstream *cmr* mutation (upstream *cmr* variant) in medium without CP, NFLX or KM. Growth rates were estimated from the exponential phase of each growth curve. Bars indicate the mean, and error bars indicate s.d. across n = 8 biological replicates. The engineered upstream *cmr* mutant did not show a detectable reduction in growth rate under these conditions ($P = 0.80$, two-sided *t*-test; n.s., not significant). **b,** Re-analysis of the Maeda et al. (2020) dataset showing the $\log_{10}$ fold change in *cmr* gene expression in lineages with (blue) or without (black) the upstream *cmr* variant 746,309 (TAATGTAA)$_{1\to2}$. In each of the chloramphenicol and puromycin experiments, one of the four evolved lineages carried this variant and three did not; the variant-carrying lineage showed an approximately eightfold higher *cmr* expression than the non-carriers. Because only one variant-carrying lineage was available per drug condition, this comparison is descriptive and no statistical test was applied. Source data are provided with this paper.

| Name | Sequence (5′→3′) | Purpose |
| --- | --- | --- |
| ybjG_cmr_int-mut-MAGE-5S4 | TTTTGCATGCAATTTCTTCGCCAATAATAATCGCGCAGAGTTTAAcAAAAGCGCAGCTAACGAGAAAGCGAATTTTGTAGCTGAAACCAC | MAGE oligonucleotide introducing 746,368 A>G in the *ybjG–cmr* intergenic region |
| ybjG_cmr_int_PCR_Fw | GCACCACGGTAATCAAATCTTTAGC | Forward primer for Sanger confirmation of 746,368 |
| ybjG_cmr_int_PCR_Rv | TGATCTTGATACAAACCGCCTCTTC | Reverse primer for Sanger confirmation of 746,368 |

**Supplementary Table 1** | Oligonucleotides used in this study. The oligonucleotide used to introduce the 746,368 A>G variant of the *ybjG–cmr* intergenic region into the unevolved *E. coli* K-12 MDS42 background by multiplex automated genome engineering (MAGE; Wang et al. 2009), and the primers used to confirm the resulting genotype by Sanger sequencing.

**Supplementary data legends**

**Supplementary Data 1 |** Detected and excluded variants. Sheet 1 lists every variant detected in the 80 sequenced samples, one row per variant per lineage, with the day-40 and day-70 calls. Sheet 2 lists the 27 variants excluded as already present in the ancestral background or fixed in the drug-free control lineages, with the reason for each.

**Supplementary notes**

**Supplementary Note 1** | Read-level detection of *ybjG–cmr* variants at day 40. Consensus genotyping called a *ybjG–cmr* variant in one of the 36 sequenced lineages at day 40. To assess whether variants were present below the consensus threshold, allele counts across the intergenic region were tabulated from the same alignments with samtools, without base alignment quality recalculation, which otherwise down-weights substitutions adjacent to the 8 bp repeat at 746,309. Depth at the six positions mutated at day 70 ranged from 23 to 134 across the 36 lineages (median 49), giving a detection limit of approximately 5%; a variant was scored as detected at a frequency of 0.10 or above. On this basis a variant was detected at day 40 in three of the nine lineages that carried one at day 70:

| Lineage | Position (AP012306) | Day-70 variant | Day-40 depth | Day-40 frequency |
|---|---|---|---|---|
| ALL→ALL 08 | 746,376 | C>A | 32 | 0.97 |
| ALL→ALL 08 | 746,396 | A>G | 32 | 0.00 |
| KM→ALL 10 | 746,394 | C>T | 43 | 0.79 |
| KM→ALL 10 | 746,349 | T>G | 58 | 0.00 |
| ALL→ALL 11 | 746,368 | A>G | 38 | 0.42 |

In both lineages carrying two substitutions at day 70, only one member of the pair was present at day 40, indicating that these haplotypes assembled sequentially. No other lineage exceeded a frequency of 0.031 at any of the six positions.
